# Altered specificity of 15-LOX-1 in the biosynthesis of 7S,14S-diHDHA implicates 15-LOX-2 in biosynthesis of resolvin D5

**DOI:** 10.1101/2020.03.25.008573

**Authors:** Steven C. Perry, Chakrapani Kalyanaraman, Benjamin E. Tourdot, William S. Conrad, Oluwayomi Akinkugbe, John Cody Freedman, Michael Holinstat, Matthew P. Jacobson, Theodore R. Holman

## Abstract

The oxylipins, 7S,14S-diHDHA and 7S,17S-diHDHA (RvD5), have been found in macrophages exudates and are believed to function as specialized pro-resolving mediators (SPM’s). Their biosynthesis is thought to proceed through sequential oxidations of docosahexaenoic acid (DHA) by lipoxygenase enzymes, specifically by h5-LOX first to 7S-HDHA, followed by h12-LOX to form 7S,14S-diHDHA or h15-LOX-1 to form 7S,17S-diHDHA (RvD5). In this work, we determined that oxidation of 7S-HpDHA to 7S,14S-diHDHA can be performed by either h12-LOX or h15-LOX-1, with similar kinetics. The oxidation at C14 of DHA by h12-LOX was expected, but the non-canonical reaction of h15-LOX-1 to make primarily 7S,14S-diHDHA was unexpected. Computer modeling suggests the alcohol on C7 of 7S-HDHA hydrogen bonds with the backbone carbonyl of I399, forcing the hydrogen abstraction from C12 to oxygenate on C14, and not C17. This result raised questions regarding synthesis of 7S,17S-diHDHA (RvD5). Strikingly, we find h15-LOX-2 oxygenates 7S-HDHA almost exclusively at C17, forming RvD5 with faster kinetics than h15-LOX-1. The presence of h15-LOX-2 in neutrophils and macrophages, suggests it may have a greater role in biosynthesizing SPM’s than previously thought. We also determined that the reactions of h5-LOX with 14S-HpDHA and 17S-HpDHA are kinetically slow compared to DHA, suggesting these may be minor biosynthetic routes *in-vivo*. Additionally, we show that 7S,14S-diHDHA and RvD5 have anti-aggregation properties with platelets at low micro-molar potencies, which could directly regulate clot resolution.

## INTRODUCTION

Inflammation plays an essential role in protecting the body from tissue injury and foreign pathogens. In its acute form, it involves the up-regulation of localized inflammatory signaling molecules and cytokines that attract neutrophils to an area of injury (1). Over time, an area of injury begins producing specialized pro-resolving mediators (SPM’s), which actively down-regulate the immune response (2), a process referred to as the resolution of inflammation. Mis-regulation of the transition to resolution can extend the early beneficial effects of acute inflammation into the damaging effects of chronic inflammation. This can contribute to cardiovascular disease (3–5), diabetes (6, 7) and autoimmune disorders (8), to name a few examples.

SPM’s are oxylipins produced by the oxygenation of fatty acids by cyclooxygenases, lipoxygenases and cytochrome P450s (9). Humans have multiple non-heme, iron lipoxygenases that catalyze the stereospecific peroxidation of polyunsaturated fatty acids containing a bis-allylic carbon moiety (10). The focus of this work are the two 15-LOX isozymes, human reticulocyte 15-lipoxygenase-1 (h15-LOX-1 or ALOX15), primarily expressed in reticulocytes and macrophages (11, 12), and human epithelial 15-lipoxygenase-2 (h15-LOX-2 or ALOX15b), expressed in macrophages, neutrophils, skin, hair roots and prostate (3, 13, 14). They are named for their ability to stereo-specifically oxygenate at C15 of arachidonic acid (AA)(10). h15-LOX-1 converts AA into a mixture of approximately 90% 15(S)-hydroperoxy-5Z,8Z,11Z,13E-eicosatetraenoic acid (15S-HpETE) and 10% 12(S)-hydroperoxy-5Z,8Z,10E,14Z-eicosatetraenoic acid (12S-HpETE), while h15-LOX-2 oxygenates substrates with strict regiospecificity (15), producing only 15-HpETE from AA. h15-LOX-1 is implicated in many diseases, such as inflammation, stroke, colorectal cancer, and atherosclerosis (11), while h15-LOX-2 has only recently been identified as playing a role in atherosclerosis and inflammation (4, 16).

Many SPM’s are synthesized at sites of inflammation and function in a paracrine or autocrine fashion. Maresin 1 (7(R),14(S)-dihydroxy-4Z,8E,10E,12Z,16Z,19Z docosahexaenoic acid, MaR1) is a SPM made from docosahexanoic acid (DHA) (17), and plays a role in the resolution of inflammation by reducing neutrophil chemotaxis and platelet aggregation and increasing phagocytosis in leukocytes (18). In conjunction with the detection of MaR1 in cell extracts, Serhan and co-workers discovered three analogues of MaR1, 7-epi-MaR1 (7S,14S-dihydroxy-4Z,8E,10E,12Z,16Z,19Z docosahexaenoic acid), 7(R),14(S)-dihydroxy-4Z,8E,10E,12E,16Z,19Z docosahexaenoic acid (i.e. *EEE*-MaR1), and 7(S),14(S)-dihydroxy 4Z,8E,10Z,12E,16Z,19Z docosahexaenoic acid (7S,14S-diHDHA, Figure 1) (17). These three analogues were shown to affect macrophage chemotaxis, but were less potent than MaR1. In particular, 7S,14S-diHDHA appeared to be biosynthesized in a di-oxygenation fashion without an epoxide intermediate, distinct from the other analogues. Its structure suggests that it is synthesized through two sequential LOX oxygenation reactions at C7 and C14, without the requirement of a hydrolase to open the epoxide intermediate (19–21). Given these oxygenation sites, as well as the enzymatic preferences of specific LOX isozymes, it is reasonable to assume that the biosynthesis of 7S,14S-diHDHA involves h5-LOX oxidizing DHA at C7, with h12-LOX oxidizing at C14, but it is unclear which oxidation occurs first (Scheme 1, Pathway 2).

**Figure 1.**
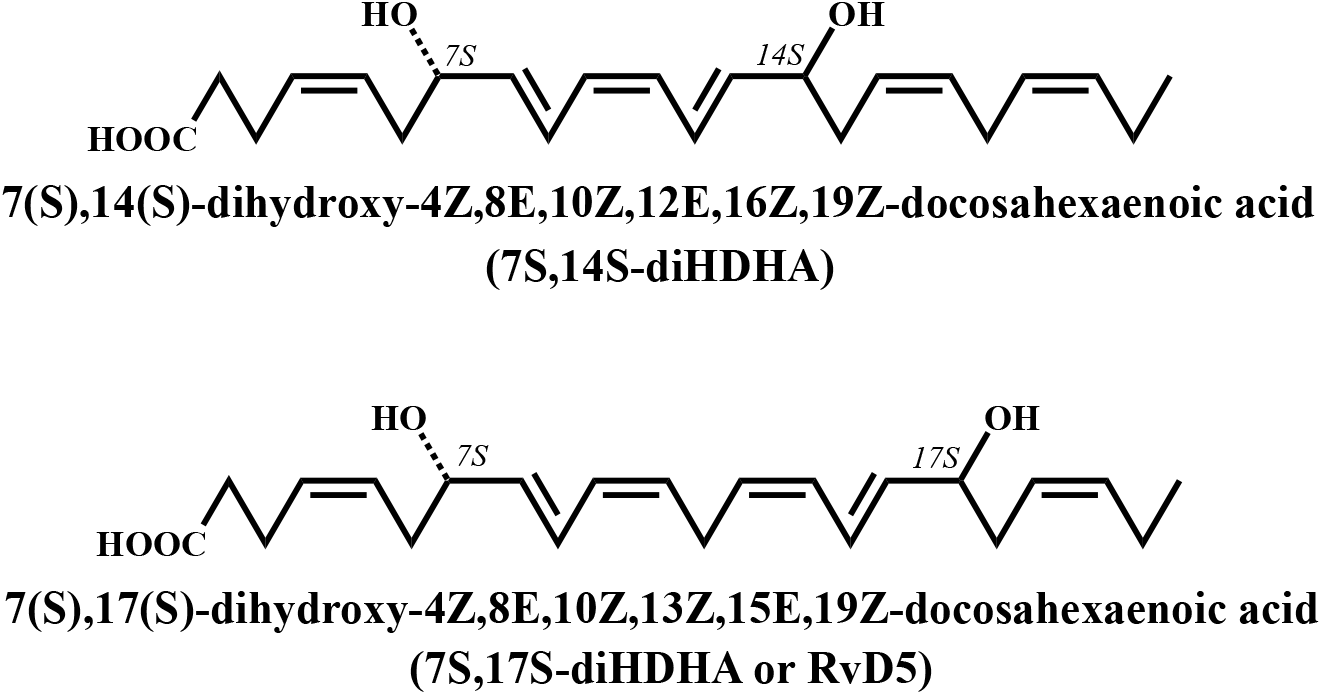
Structures of 7(S),14(S)-dihydroxy-4Z,8E,10Z,12E,16Z,19Z-docosahexaenoic acid (7S,14S-diHDHA), and 7(S),17(S)-dihydroxy-4Z,8E,10Z,13Z,15E,19Z-docosahexaenoic acid (7S,17S-diHDHA or RvD5). Note that the molecules are drawn with an artificial straight chain shape in order to highlight their similarities and differences more clearly.

**Scheme 1.**
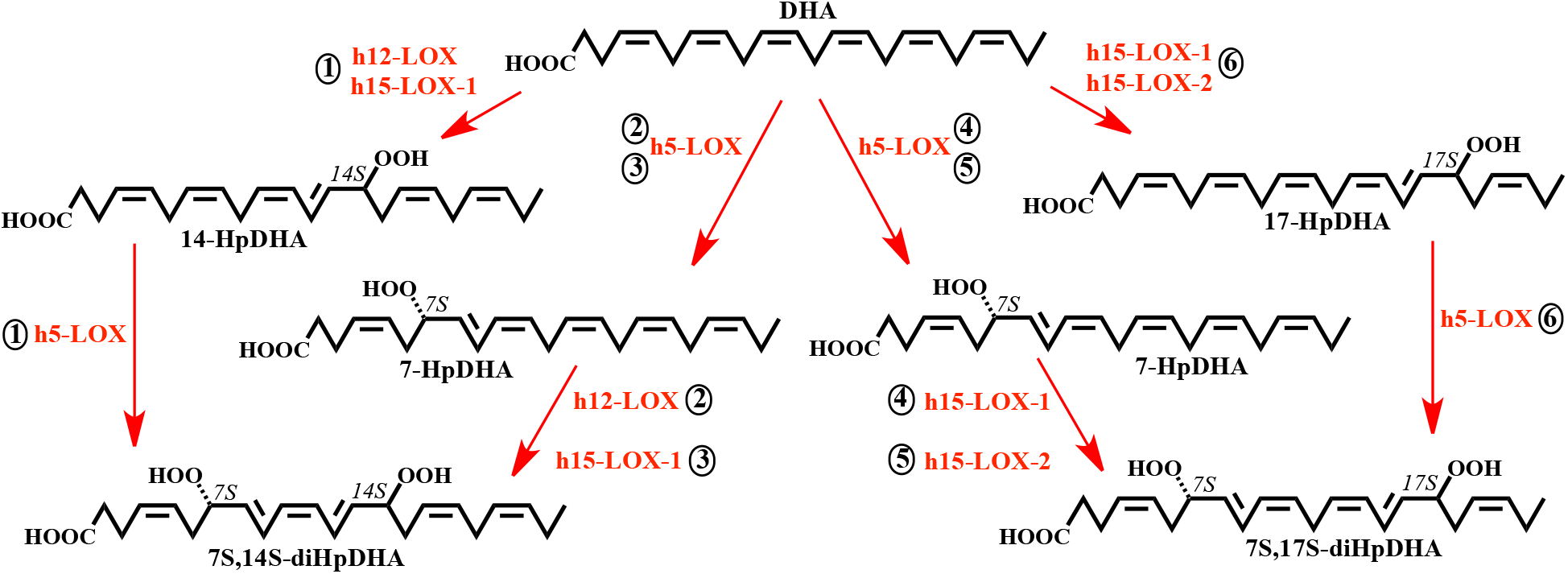
The synthesis of 7S,14S-diHDHA from DHA may occur through one of three possible pathways: reaction of DHA with h5-LOX followed by h12-LOX (2), or reaction of DHA with h5-LOX followed h15-LOX-1 (3) or reaction of DHA with h12-LOX or h15LOX-1 followed by h5-LOX (1). The synthesis of RvD5 from DHA may also occur through three possible pathways: reaction of DHA with h5-LOX followed by h15-LOX-1 (4), or reaction of DHA with h5-LOX followed h15-LOX2 (5) or reaction of DHA with h12-LOX, h15-LOX-1 or h15LOX-2 followed by h5-LOX (6).

The other SPM which is of importance with regard to this work is Resolvin D5 (RvD5 or 7(S),17(S)-dihydroxy 4Z,8E,10Z,13Z,15E,19Z docosahexaenoic acid (7S,17S-diHDHA), Figure 1. RvD5 is a D-series resolvin made from DHA (22), and has been identified in human blood, hemorrhagic exudates and synovial fluid (23, 24). RvD5 enhances phagocytosis in neutrophils and macrophages and reduces expression of the pro-inflammatory molecules, NF-κB and TNF-α (25). Like 7S,14S-diHDHA, the production of RvD5 appears to require two oxygenation events and is proposed to occur through the sequential reaction of two lipoxygenases (24, 26). Since many cell types contain only a single lipoxygenase, oxylipins with multiple oxygenation sites are often created by the interaction of multiple cell types in an area of inflammation through a process called transcellular biosynthesis (27–29). RvD5 has been isolated from neutrophils (30) and is produced late in the process of blood coagulation, increasing over the lifetime of a clot (31). PMN’s have been shown to convert 17(S)-hydroxy 4Z,7Z,10Z,13Z,15E,19Z docosahexaenoic acid (17S-HDHA) into 4(S),17(S)-dihydroxy 5E,7Z,10Z,13Z,15E,19Z docosahexaenoic acid (4S,17S-diHDHA) and RvD5 (24), suggesting that h5-LOX reacts with 17-HDHA to produce RvD5.

As stated above, LOX isozymes play a key role in the biosynthesis of SPMs, however, the biosynthetic molecular mechanism for specific SPMs is poorly defined. For 7S,14S-diHDHA, the current understanding of its biosynthetic pathway is based on the substrate specificity of h5-LOX and h12-LOX with AA, and experimental evidence that the non-specific LOX inhibitor, baicalein, reduces the production of MaR1 and 14(S)-dihydroxy-4Z,7Z,10Z,12E,16Z,19Z-docosahexaenoic acid (14S-HDHA) (32). However, baicalein is not a selective inhibitor (33) and it is well recognized that LOXs can produce different products depending on the fatty acids or oxylipin used (34). For example, h5-LOX produces primarily 5(S)-hydroperoxy-6E,8Z,11Z,14Z-eicosatetraenoic acid (5S-HpETE) from AA, but it produces multiple products when DHA is the substrate (35), presumably due to the length and unsaturation differences between AA and DHA, which presumably affects substrate positioning for hydrogen atom abstraction. LOX’s also vary widely in their ability to tolerate oxygenated substrates, as reflected in their kinetic parameters (19). In addition, the specific hydrogen atom abstracted can be affected by the nature of the substrate (34). h5-LOX abstracts a hydrogen atom from C7 of AA to produce 5S-HpETE, but it can subsequently abstract from C10 from 5S-HpETE to produce the 5,6-epoxide (36). This variability in product distribution is also seen with h15-LOX-1, where approximately 10% of the product made from AA is 12S-HpETE, the non-canonical product. Kuhn and coworkers proposed that this lack of specificity is linked to human evolution and specifically the human inflammatory response, with only higher primates having this LOX function (37). This ability of h15-LOX-1 to oxygenate either C15 or C12 on AA raises the possibility that h15-LOX-1 could generate 7S,14S-diHDHA from DHA (38), as shown in Scheme 1, Pathway 3.

For RvD5, the generally accepted biosynthetic pathway is through transcellular biosynthesis, but a specific biosynthetic sequence has not been clearly identified. One proposed pathway is that h5-LOX, expressed in macrophages (39, 40) and PMN’s (24), produces 7(S)-hydroxy-4Z,8E,10Z,13Z,16Z,19Z-docosahexaenoic acid (7S-HDHA) from DHA, and then h15-LOX-1 reacts with 7S-HDHA to produce RvD5 by oxidation at C17. This is a reasonable assumption since h15-LOX-1 oxygenates C17 of DHA efficiently; however, h15-LOX-2 is expressed at higher levels than h15-LOX-1 in both neutrophils and macrophages (4, 13, 41), so it is possible that h15-LOX-2, and not h15-LOX-1, may be involved in the formation of RvD5 from 7S-HDHA (Scheme 1). It should be noted that LOXs produce the hydroperoxy product, but the reducing environment of the cell quickly reduces them to the hydroxy products (42), such as 7S,14S-diHDHA and 7S,17S-diHDHA (RvD5).

Given the biological importance of SPMs, such as 7S,14S-diHDHA and 7S,17S-diHDHA (RvD5), and the multiple possible biosynthetic pathways for their production, we investigated two critical aspects of the biosynthetic routes to producing these two SPMs: the ease of which LOXs can perform the proposed reactions *in vitro* (i.e. kinetics), and the specificity of LOXs when the substrate is an oxylipin, as opposed to the un-oxidized fatty acid. We probed these effects in the production of both 7S,14S-diHDHA and 7S,17S-diHDHA (RvD5) and determined that not only are the rates greatly affected by the nature of the oxylipin substrate, but also the product profile is altered, raising the possibility that unrecognized, non-canonical LOXs may be involved in SPM production.

## MATERIALS AND METHODS

### Expression and Purification of h15-LOX-1, h15-LOX-2, h12-LOX, and h5-LOX

Overexpression and purification of wild-type h15-LOX-1 (Uniprot entry P16050), h12-LOX (Uniprot entry P18054), h5-LOX (Uniprot entry P09917) and h15-LOX-2 (Uniprot entry O15296) were performed as previously described (78–80). The purity of h15-LOX-1 and h12-LOX were assessed by SDS gel to be greater than 85%, and metal content was assessed on a Finnigan inductively-coupled plasma-mass spectrometer (ICP-MS), via comparison with iron standard solution. Cobalt-EDTA was used as an internal standard. The wt h5-LOX used in the kinetics of this work was not purified due to a dramatic loss in activity and was therefore prepared as an ammonium sulfate-precipitate. The amount of h5-LOX contained in the ammonium-sulfate pellet was assessed by comparing values obtained by a Bradford assay and quantitative western blotting using purified stable-h5-LOX mutant as a positive control. Western blots were performed using rabbit anti-5-lipoxygenase polyclonal (Cayman chemicals) primary antibody diluted 1:500 and goat anti-rabbit-HRP (Abcam) secondary antibody diluted 1:5000. The stable-h5-LOX mutant was expressed in Rosetta 2 cells (Novagen) transformed with the pET14b-Stable-5-LOX plasmid (a gift from Marcia Newcomer of Louisiana State University) and grown in Terrific Broth containing 34 µg mL^-1^ chloramphenicol and 100 µg mL^-1^ ampicillin at 37 °C for 3.5 h and then placed at 20 °C for an additional 26 h. Cells were pelleted and resuspended in 50 mM Tris (pH 8.0), 500 mM NaCl, 20 mM imidazole with 1 µM Pepstatin, 100 µM PMSF, and DNaseI (2 Kunitz/g) (Sigma). The cells were lysed in a French pressure cell and centrifuged at 40,000xg for 20min. Lysate was applied to a Talon nickel-IDA Sepharose column and eluted with 50 mM Tris pH 8.0, 500 mM NaCl, 200 mM imidazole. The final product was stored at −80 °C with 10% glycerol.

### Production and isolation of oxylipins

7(S)-hydroperoxy-4Z,8E,10Z,13Z,16Z,19Z-docosahexaenoic acid (7S-HpDHA) was synthesized by reaction of DHA (25-50 µM) with h5-LOX. The reaction was carried out for 2 hours in 800 mL of 25mM HEPES, pH 7.5 containing 50 mM NaCl, 100 µM EDTA and 200 µM ATP. The reaction was quenched with 0.5% glacial acetic acid, extracted 3 times with 1/3 volume dichloromethane and evaporated to dryness under N2. The Products were purified isocratically via high performance liquid chromatography (HPLC) on a Higgins Haisil Semi-preparative (5 µm, 250 mm x 10 mm) C18 column with 50:50 of 99.9% acetonitrile, 0.1% acetic acid and 99.9% water, 0.1% acetic acid. 7(S)-hydroxy-4Z,8E,10Z,13Z,16Z,19Z-docosahexaenoic acid (7S-HDHA) was synthesized as performed for 7S-HpDHA with trimethylphosphite added as a reductant prior to HPLC. 7S,14S-diHDHA was synthesized by reaction of 20 µM 7S-HDHA with h15-LOX-1 in 300 mL of 25mM HEPES, pH 7.5 and purified as described above. 14(S)-hydroperoxy 4Z,7Z,10Z,12E,16Z,19Z-docosahexaenoic acid (14S-HpDHA) was synthesized by reaction of DHA (25-50 µM) with h12-LOX. The reaction was carried out for 30 minutes in 1000 mL of 25 mM HEPES, pH 8.0 The reaction was quenched with 0.5% glacial acetic acid, extracted 3 times with 1/3 volume dichloromethane and evaporated to dryness under N2. The Products were purified isocratically via HPLC on a Phenomenex Luna semi-preparative (5µm, 250mm x 10mm) silica column with 99:1 of 99.9% hexane, 0.1% trifluoroacetic acid and 99.9% isopropanol, 0.1% trifluoroacetic acid. 14(S)-hydroxy 4Z,7Z,10Z,12E,16Z,19Z-docosahexaenoic acid (14S-HDHA) was synthesized as performed for 14S-HpDHA with trimethylphosphite added as a reductant prior to HPLC. 17(S)-hydroperoxy 4Z,7Z,10Z,13Z,15E,19Z-docosahexaenoic acid (17S-HpDHA) was synthesized by reaction of DHA (25-50 µM) with soybean 15-LOX. The reaction was carried out for 1 hour in 800 mL of 100 mM Borate, pH 9.0. 17S-HDHA was synthesized by reducing 17S-HpDHA with trimethylphosphite, prior to HPLC. 7(S),17(S)-dihydroxy docosahexaenoic acid (7S,17S-diHDHA or RvD5) was synthesized by reacting 7S-HDHA (20 µM) with h15-LOX-2 in 300 mL of 25 mM HEPES, pH 7.5, with subsequent reduction/purification, as described above. The isolated products were assessed to be greater than 95% pure by LC-MS/MS.

### Steady State Kinetics of h15-LOX-1 and h12-LOX with DHA, 7S-HDHA and 7S-HpDHA

h15-LOX-1 reactions were performed, at ambient temperature, in a 1 cm^2^ quartz cuvette containing 2 mL of 25 mM HEPES, pH 7.5 with substrate (DHA, 7S-HDHA or 7S-HpDHA). DHA concentrations were varied from 0.25-10 µM, 7S-HDHA concentrations were varied from 0.3-15 µM and 7S-HpDHA concentrations were varied from 0.3-20 µM. Concentration of DHA was determined by measuring the amount of 17S-HpDHA produced from complete reaction with soybean lipoxygenase-1 (sLO-1). Concentrations of 7S-HDHA and 7S-HpDHA were determined by measuring the absorbance at 234 nm. Reactions were initiated by the addition ∼200 nM h15-LOX-1 and were monitored on a Perkin-Elmer Lambda 45 UV/VIS spectrophotometer. Product formation was determined by the increase in absorbance at 234 nm for 7S-HpDHA (ε_234nm_ = 25,000 M^-1^ cm^-1^) and 270 nm for 7S,14S-diHDHA (ε_270nm_ = 40,000 M^-1^ cm^-1^)(32, 81, 82). 7S,17S-diHDHA has an absorbance max of 245 nm, however, due to overlap with the substrate peak at 234 nm formation of this product was measured at 254 nm using an extinction coefficient of 21,900 M^-1^ cm^-1^ to adjust for the decreased rate of absorbance change at this peak shoulder (19). KaleidaGraph (Synergy) was used to fit initial rates (at less than 20% turnover), as well as the second order derivatives (*k_cat_/K_M_*) to the Michaelis-Menten equation for the calculation of kinetic parameters. h12-LOX reactions were performed similarly as for h15-LOX-1 except that buffers were 25 mM HEPES, pH 8 and reactions were initiated by the addition ∼50 nM h12-LOX.

### Steady State Kinetics of h5-LOX with DHA, 14S-HDHA, 14S-HpDHA, 17S-HDHA and 17S-HpDHA

All h5-LOX kinetic reactions were carried out in 25mM HEPES, pH 7.5 containing 50mM NaCl, 100 µM EDTA and 200 µM ATP. Steady state kinetics of h5-LOX with DHA were determined as for h15-LOX-1. *V_max_* assay of h5-LOX with 14S-HDHA, 14S-HpDHA, 17S-HDHA, 17S-HpDHA were carried out in 15 ml of buffer, containing 10 µM substrate Reactions were initiated by the addition ∼600 nM ammonium-sulfate precipitated h5-LOX and were monitored on a Perkin-Elmer Lambda 45 UV/VIS spectrophotometer. Reactions were quenched at 0, 5, 10, 20, 30 and 60 minutes. Each quenched reaction was extracted three times with 1/3 volume of DCM and reduced with trimethylphosphite. The samples were then evaporated under a stream of N_2_ to dryness and reconstituted in 90:10 acetonitrile:water containing 0.1% formic and 3 µM 13-HODE as an internal standard. Control reactions without enzyme were also conducted and used for background subtraction, ensuring oxylipin degradation products were removed from analysis. Reactions were analyzed via LC-MS/MS using a Synergi 4 µM Hydro-RP 80 Å C18 LC-Column with a polar endcapping (150 x 2 mm). Mobile phase solvent A consisted of 99.9% water, 0.1% formic acid and solvent B consisted of 99.9% acetonitrile, 0.1% formic acid. Analysis was carried out over 60 min using isocratic 50:50 A:B for 0-30 min followed by a gradient from 50:50 A:B to 75:25 A:B, from 30-60 min. The chromatography system was coupled to a Thermo-Electron LTQ LC-MS/MS for mass analysis. All analyses were performed in negative ionization mode at the normal resolution setting. MS2 was performed in a targeted manner with a mass list containing the following m/z ratios ± 0.5: 343.4 (HDHA’s), 359.4 (diHDHA’s), and 375.4 (triHDHA’s). Products were identified by matching retention times, UV spectra, and fragmentation patterns to known standards, or in the cases were MS standards were not available, structures were deduced from comparison with known and theoretical fragments. MS/MS fragments used to identify 7,14-diHDHA included: 341, 297, 221, 177, 141 and 123. MS/MS fragments used to identify 7,17-diHDHA included: 341, 297, 261, 243, 199 and 141. MS2 integrated peak areas were normalized to a 13-HODE internal standard and fitted to determine *V_max_*. The catalytic activity relative to protein weight for h5-LOX was calculated by comparing *V_max_* to protein concentration which was determined by quantitative western, described above. For comparison of biosynthetic flux with h5-LOX, *K_cat_* values for each enzyme and substrate were converted to V_max_ at 10 µM using the Michaelis-Menten equation and then multiplied by the percentage of total product represented by each reaction product.

### Steady State Kinetics of h15-LOX-2 with DHA, 7S-HDHA and 7S-HpDHA

h15-LOX-2 reactions were performed, at room temperature, in a 1 cm^2^ quartz cuvette containing 2 mL of 25 mM HEPES, pH 7.5 with substrate (DHA, 7S-HDHA and 7S-HpDHA.) DHA concentrations were varied from 0.75-50 µM, 7S-HDHA concentrations were varied from 0.6-30 µM, 7S-HpDHA concentrations were varied from 0.6-25 µM. The concentration of DHA was determined by measuring the amount of 17S-HpDHA produced from complete reaction with soybean lipoxygenase-1 (sLO-1) and confirmed by measuring turnover with h15-LOX-2. Concentrations of 7S-HDHA and 7S-HpDHA were determined by measuring absorbance at 234 nm. Reactions were initiated by the addition h15-LOX-2 (∼220 nM final concentration) and were monitored on a Perkin-Elmer Lambda 45 UV/VIS spectrophotometer. Product formation was determined by the change in absorbance at 234 nm for 7S-HDHA (ε_234nm_ = 25,000 M^-1^ cm^-1^), 270 nm for 7S,14S-diHDHA (ε_270nm_ = 40,000 M^-1^ cm^-1^), and 254 nm for 7S,17S-diHDHA (ε_254nm_ = 21,900 M^-1^ cm^-1^) (19, 32, 81, 82),. KaleidaGraph (Synergy) was used to fit initial rates (at less than 20% turnover), as well as the second order derivatives (*k_cat_/K_M_*) to the Michaelis-Menten equation for the calculation of kinetic parameters.

### Product Analysis of LOX reactions

Reactions were carried out in 2 mL of 25 mM HEPES, pH 7.5 with stirring at ambient temperature. Reactions with DHA and AA contained 10 µM substrate and ∼200 nM h15-LOX-1 and h15-LOX-2 in 2 ml of buffer. Those with h15-LOX-1 and 7S-HDHA, 7S-HpDHA, 14S-HDHA and 14S-HpDHA contained 20 µM substrate and ∼600 nM of enzyme in 4 ml of buffer. Those with, h15-LOX-2 and 7S-HDHA, 7S-HpDHA, 17S-HDHA and 17S-HDHA contained 4 mL of buffer, 10 µM substrate and ∼450 nM of enzyme. The reaction of h5-LOX with DHA contained 10 µM substrate and ∼400 nM ammonium sulfate-precipitated enzyme and was analyzed by LCMS, as described above for h5-LOX kinetics. Reactions of h15-LOX-1, h15-LOX-2 and h12-LOX were analyzed by LCMS as described above, with the exception that the LC gradient was carried out over 25 min using isocratic 40:60 A:B for 0-10 min followed by a gradient from 40:60 A:B to 75:25 A:B, from 10-25 min. In order to minimize the formation of secondary products, reactions were quenched at less than 50% turnover. The presence of tri-HDHA secondary products was assessed by monitoring at MS/MS filter of 359.4. Any tri-HDHA’s observed were quantitated and included in the area of their respective parent di-HDHA.

### Chiral Chromatography

7S,14S-diHDHA and 7S,17S-diHDHA were synthesized by h15-LOX-1 and h15-LOX-2, respectively and isolated via HPLC as described above. 7S,14S-diHDHA produced by h5-LOX was synthesized in 15 mL of buffer containing 10 µM 14S-HDHA and ∼6 µM of ammonium-sulfate precipitated h5-LOX. Purified 7S,14S-diHDHA and 7S,17S-diHDHA were analyzed via LC-MS/MS using a Chirapak AD-RH 2.1 x 150 mm 5 µM chiral column coupled to a Thermo electron LTQ. Mobile phase solvent A consisted of 99.9% water, 0.1% formic acid and solvent B consisted of 99.9% acetonitrile, 0.1% formic acid. Analysis was carried out over 60 min using isocratic 35:65 A:B for 0-30 min followed by a gradient from 35:65 A:B to 75:25 A:B, from 30-60 min. MS/MS conditions were the same as described above. Retention times and fragmentations were compared to RvD5, Mar1 and 7-epi-Mar standards purchased from Cayman Chemicals (Ann Arbor, MI).

### Oxylipin titration into human platelets

The University of Michigan Institutional Review Board approved all research involving human volunteers. Washed platelets were isolated from human whole blood via serial centrifugation and adjusted to 3.0 x 10^8^ platelets/mL in Tyrode’s buffer (10 mM HEPES, 12 mM NaHCO_3_, 127 mM NaCl, 5 mM KCl, 0.5 mM NaH_2_PO_4_, 1 mM MgCl_2_, and 5 mM glucose), as previously published. Platelets (250 µL at 3.0 x10^8^ platelets/mL) were dispensed into glass cuvettes and incubated with the indicated oxylipin in half-log increments (0-10 µM) for 10 minutes at 37°C. Oxylipin-treated platelets were stimulated with 0.25 µg/mL of collagen (Chrono-log), under stirring conditions (1100 rpm) at 37°C, in a Chrono-log Model 700D lumi-aggregometer and platelet aggregation was recorded for six minutes.

In order to determine if 7S-HDHA was enzymatically converted to another chemical *ex vivo*, one mL of platelets (1.0 x10^9^ platelets/mL) was incubated with either 10 µM 7S-HDHA, 10 µM 5S-HETE (positive control) or vehicle (DMSO) for 10 minutes at 37° C and then pelleted by centrifugation at 1000g for 2 minute. Supernatant was transferred to a fresh tube and snap frozen. Oxylipins were extracted and analyzed via UPLC-MS/MS, as described previously,*^9^* The m/z transitions for 7S-HDHA, 7S,14S-diHDHA, 7S,17S-diHDHA were 343.5à141, 359.5à141, 359.5à199, respectively.

### Molecular Modeling

A structure of h15-LOX-1 (ALOX15) is not available in the Protein Data Bank (PDB). Therefore, we constructed a homology model of h15-LOX-1 from its sequence (Uniprot ID: P16050) using the substrate mimetic inhibitor bound, high-resolution structure of porcine 12-LOX (PDB ID: 3rde). The h15-LOX-1 sequence is 86% identical to the porcine 12-LOX sequence, with both being considered ALOX15 genes. The homology model was constructed using Prime software (version 5.4, Schrodinger Inc). During the modeling step we retained the co-crystallized inhibitor, metal ion (Fe^3+^), and a hydroxide ion that coordinated the metal ion from the homolog structure. The model was subsequently energy minimized using Protein Preparation Wizard module of Maestro (version 11.8, Schrodinger Inc). During this step, hydrogen atoms were added to the protein, co-crystallized ligand and the hydroxide ion. Hydrogen atoms of the titratable residues were adjusted and side chains of Tyr, Thr, Ser, Asn and Gln were optimized so they could make better hydrogen bonding interactions. The structure was finally minimized such that the heavy atoms do not move beyond 0.3Å from their starting positions. Structures of DHA and 7S-HDHA were modeled using Edit/Build panel of Maestro and energy minimized using LigPrep software (Schrodinger Suite 2018-4, Schrodinger Inc). Both DHA and 7S-HDHA were docked to the h15-LOX-1 model using the docking software Glide (version 8.1, Schrodinger Inc) with the extra-precision (XP) docking score. We used the co-crystallized ligand coordinates to define the binding pocket. During the ligand-docking step, the protein was initially kept rigid, but despite extensive ligand conformation sampling, no low-energy docking pose was identified for the ligands. Therefore, we used InducedFit docking (Schrodinger Inc), in which both protein and ligand are treated flexibly. Specifically, during the initial docking step, side chains of Phe352, Ile413, Ile417, Met418, Cys559, and Leu588 were deleted to accommodate DHA and 7S-HDHA, because they appeared to block key portions of the active site. After generating the initial docking pose, side chain rotamers of these residues plus Arg402 and Gln595 (which have the potential to form hydrogen bonds with the ligand) were optimized. These docking and side chain optimization steps were iterated until a converged low energy docking pose was obtained.

For h15-LOX-2, structure-based docking calculations were performed using the available crystal structure of human ALOX15b (PDB id: 4nre). From the crystal structure, we retained the protein, Fe^3+^ ion, co-crystallized inhibitor, polyoxyethylene detergent (C8E), present in the active site and a hydroxide ion that co-ordinates the metal ion, all other atoms were deleted. Prior to docking, the structure was subjected to the protein-preparation step using Maestro’s Protein-Preparation Wizard (Maestro version 11.8, Schrodinger Inc). Although the co-crystallized inhibitor bound in a U-shaped binding mode, it did not have a carboxylate group, and it had only 21 atoms in the main chain as opposed to 22 atoms present in DHA or 7S-HDHA. Therefore, we performed flexible-receptor flexible-ligand docking using InducedFit docking software (Schrodinger Inc). During induced-fit docking only the following active site residues were treated flexibly: Phe365, Val426, Val427, Arg329 and Asp602.

## RESULTS

### Biosynthesis of 7S,14S-diHDHA

As discussed in the introduction, there are three proposed routes by which 7S,14S-diHDHA can be synthesized by LOXs (Scheme 1, Pathways 1-3). These three pathways were therefore investigated *in vitro* to implicate possible *in vivo* biosynthetic routes. It should be emphasized that since these are di-oxygenation pathways, the di-hydroperoxide is formed in each of the pathways, however the biologically isolated molecule, 7S,14S-diHDHA, is the reduced form due to cellular glutathione peroxidases.

### Pathway 1: DHA with h12-LOX to 14S-HpDHA, then h5-LOX to 7S,14S-diHpDHA

The first possible route for biosynthesis of 7S,14S-diHDHA involves oxygenation of DHA by h12-LOX to form 14S-HpDHA (Scheme 1, Pathway 1). In order to study the ability of h12-LOX to form 14S-HpDHA, steady state kinetics and product profiles were measured. The *k_cat_* for h12-LOX with DHA was measured to be 14 sec^-1^, while the *k_cat_/K_M_* was found to be 13 sec^-1^µM^-1^ (Table 1), demonstrating that DHA is a comparable substrate to AA for h12-LOX *in vitro* (43, 44). The product profile indicates that the majority of oxylipin is oxygenated at C14 (greater than 80%), with minor amounts oxygenated at C11, as previously reported (45, 46).

**Table 1.** Steady-state kinetic values of h12-LOX and h15-LOX-1 with DHA, 7S-HDHA and 7S-HpDHA.

| Enzyme | Substrate | $k_{cat}$ (sec<sup>-1</sup>) | $k_{cat}/K_M$<br>(sec<sup>-1</sup>μM<sup>-1</sup>) | $K_M$ (μM) |
| --- | --- | --- | --- | --- |
| h12-LOX | AA | 9.7 ± 1 | 5.8 ± 0.4 | 1.7 ± 0.3 |
|  | DHA | 14 ± 1 | 13 ± 2 | 1.1 ± 0.2 |
|  | 7S-HDHA | 3.3 ± 0.2 | 1.8 ± 0.5 | 1.8 ± 0.5 |
|  | 7S-HpDHA | 0.68 ± 0.1 | 0.25 ± 0.1 | 2.7 ± 1 |
| h15-LOX-1 | AA | 10 ± 1 | 2.1 ± 0.3 | 5.0 ± 0.3 |
|  | DHA | 0.95 ± 0.1 | 0.68 ± 0.2 | 1.4 ± 0.3 |
|  | 7S-HDHA | 3.1 ± 0.1 | 0.51 ± 0.09 | 6.1 ± 0.5 |
|  | 7S-HpDHA | 0.57 ± 0.08 | 0.080 ± 0.02 | 6.9 ± 0.8 |

After DHA reacts with h12-LOX to form 14S-HpDHA, it may subsequently react with h5-LOX to form 7S,14S-diHpDHA (Scheme 1, Pathway 1), the oxidized form of 7S,14S-diHDHA. In order to study the ability of h5-LOX to react with 14S-HDHA and 14S-HpDHA, steady-state kinetic values and product profiles were assessed but no activity was observed by UV-vis spectroscopy (i.e. no change in absorbance at 254, 270 or 302 nm being detected). Using the more sensitive LC-MS/MS method, the *V_max_* values for 14S-HDHA and 14S-HpDHA were determined to be 0.00038 and 0.0015 (mol/sec^-1^mol^-1^) at 10 µM substrate, respectively (Table 2). If we compare the *V_max_* at 10 µM of DHA (*vide supra*), the *V_max_* values of 14S-HDHA and 14S-HpDHA are 368-fold and 93-fold slower than that of DHA, indicating that 14-oxylipins are poor substrates of h5-LOX. Even with this lowered kinetic rate, the major product in both reactions was the 7S,14S-oxylipin, along with trace amounts of 8,14-oxylipin and tri-oxygenated products.

**Table 2.** V_max_ values of h5-LOX, h12-LOX and h15-LOX-1 with DHA, 14S-HDHA, 14S-HpDHA, 7S-HDHA and 7S-HpDHA, determined at 10 µM substrate concentration. The *V_max_* for h5-LOX was approximated by determining the amount of h5-LOX in the ammonium sulfate pellet by Western analysis relative to a known standard. *Biosynthetic flux is calculated by multiplying each *V_max_* by the percentage of total product from that reaction that serves as substrate for the next step towards synthesis of 7S,14S-diHDHA. Reactions with DHA represent the first of two biosynthetic steps, while reactions with oxylipins represent the second.

| Enzyme | Substrate | $V_{max}$<br>mol/sec<sup>-1</sup>mol<sup>-1</sup> | 7S,14S-diHDHA<br>biosynthetic<br>pathway flux* |
| --- | --- | --- | --- |
| h5-LOX | DHA | 0.14 | 0.073 |
|  | 14S-HDHA | 0.00038 | 0.00038 |
|  | 14S-HpDHA | 0.0015 | 0.0015 |
| h12-LOX | DHA | 12 | 9 |
|  | 7S-HDHA | 2.8 | 2.3 |
|  | 7S-HpDHA | 0.53 | 0.44 |
| h15-LOX-1 | DHA | 0.84 | 0.17 |
|  | 7S-HDHA | 1.9 | 1.7 |
|  | 7S-HpDHA | 0.30 | 0.19 |

### Pathway 2: DHA and h5-LOX to 7S-HpDHA, then h12-LOX to 7S,14S-diHpDHA (oxidized form of 7S,14S-diHDHA)

The second possible route for biosynthesis of 7S,14S-diHDHA begins with oxygenation of DHA by h5-LOX to form 7S-HpDHA (Scheme 1, Pathways 2 and 3). The ability of h5-LOX to form products from DHA was assessed through kinetic measurements and product profiles. The *V_max_* for DHA at 10 µM was determined to be 0.14 mol/sec^-1^mol^-1^ (Table 2), similar to the reported value of 0.14 mol/sec^-1^mol^-1^ for AA (converted from *k*_cat_ and *K*_M_ at 10 µM AA in reference (34)). Interestingly, the reaction of h5-LOX with DHA was found to produce 8 different products, with 7S-HpDHA being the major product at 52%, and 7 minor products comprising the remaining 48% (Table 3 and Supporting Information, Figure 1S). The 4-product was not observed, consistent with results obtained from h5-LOX in neutrophils (35, 47) and monocytes (48). It is important to note that the increased non-specificity of h5-LOX with DHA relative to AA is a common observation for other LOX isozymes as well (*vide infra*). This appears to be a function of the structural difference between DHA and AA, with the increased length and unsaturation of DHA leading to a substrate-binding mode that promotes greater product diversity relative to AA.

**Table 3.** Distribution of products created by reaction of h5-LOX with DHA. 7S-HpDHA is the major product, at 52% of the total and 7 minor products comprise the remaining 48%.

| Substrate | 7-HpDHA | 8-HpDHA | 11<br>HpDHA | 13-HpDHA | 14-HpDHA | 16-HpDHA | 17-HpDHA | 20-HpDHA |
| --- | --- | --- | --- | --- | --- | --- | --- | --- |
| DHA | 52 ± 9% | 1 ± 1% | 4 ± 2% | 2 ± 1% | 18 ± 4% | 4 ± 2% | 16 ± 6% | 3 ± 2% |

The 7S-HpDHA produced by h5-LOX from DHA contains bisallylic carbons that may react further with h12-LOX to produce the dioxygenated oxylipin, 7S,14S-diHDHA (Scheme 1, Pathway 2). To test this possibility, steady-state kinetic values were determined for the reaction of h12-LOX with 7S-HDHA and 7S-HpDHA. The *k_cat_* for 7S-HDHA and 7S-HpDHA were found to be 3.3 sec^-1^ and 0.68 sec^-1^, respectively and the *k_cat_/K_M_*’s for 7S-HDHA and 7S-HpDHA were found to be 1.8 sec^-1^µM^-1^ and 0.25 sec^-1^µM^-1^, respectively (Table 1), which are markedly greater than the kinetic parameters of h5-LOX with the 14-oxylipins. The products of h12-LOX reacting with 7S-HDHA were exclusively dioxygenated oxylipins, with 82% of 7S,14S-diHDHA and 18% of 7S,17S-diHDHA being generated (Table 4 and Supporting Information, Figure 2S). The production of 7S,17S-diHDHA is a remarkable result since h12-LOX is known to produce mostly 14S-HpDHA and only a minor amount of 11-HpDHA from DHA. The formation of 11-HpDHA indicates that DHA inserts deeper into the cavity for hydrogen atom abstraction at C9. However, with 7S-HDHA as the substrate, the minor product is 7S,17S-diHDHA, not 7S,11S-diHDHA, suggesting that 7S-HDHA does not enter as deep into the active site, allowing for abstraction at C15 and the generation of 7S,17S-diHDHA. The reaction of 7S-HpDHA with h12-LOX produced comparable products as that from 7S-HDHA, indicating that no dehydration occurred to form the epoxide or its derivatives (Table 4).

**Table 4.** Distribution of products produced by h12-LOX from 7S-HDHA as determined by LCMS/MS. The major product of the reaction of h12-LOX with 7S-HDHA is 7S,14S-diHpDHA. Note, the abbreviation, “14-product” etc., is used to simplify the complexity of the table labels. *A peak of approximately 3% the total area was observed that had a parent mass of a di-HDHA, however its exact nature was undetermined due to its small area.

| Substrate | 14-product | 17-product |
| --- | --- | --- |
| 7S-HDHA | 82 ± 6% | 18 ± 4% |
| 7S-HpDHA | 79* ± 4% | 18 ± 3% |

### Pathway 3: DHA and h5-LOX to 7S-HpDHA, then h15-LOX-1 to 7S,14S-diHpDHA (oxidized form of 7S,14S-diHDHA)

The third pathway involves the non-canonical reaction of h15-LOX-1 with 7S-HpDHA to produce 7S,14S-diHpDHA, which is subsequently reduced to 7S,14S-diHDHA (Scheme 1, Pathway 3). h15-LOX-1 was reacted with DHA, 7S-HDHA and 7S-HpDHA, with 7S-HDHA demonstrating a comparable rate to that of DHA, but 7S-HpDHA showing a slower rate, with *k_cat_/K_M_* values of 0.68, 0.51 and 0.08 sec^-1^µM^-1^ for DHA, 7S-HDHA and 7S-HpDHA, respectively (Table 1). Even more surprising, the *k_cat_* value for 7S-HDHA was over 3-fold greater than that for DHA. These results indicate that DHA is still the more efficient substrate at low concentration (i.e. the fastest rate of substrate capture, *k_cat_/K_M_*) (49), but 7S-HDHA is an alternative substrate to DHA at high concentration, having the fastest rate of product release (*k_cat_*) (49).

To determine the oxylipins generated by h15-LOX-1 from 7S-HDHA and 7S-HpDHA, the reaction products were isolated and analyzed via LC-MS/MS. Surprisingly, when h15-LOX-1 was reacted with 7S-HDHA, 90% of the products were 7S,14S-diHpDHA (i.e. the oxidized form of 7S,14S-diHDHA), with only 10% of 7S,17S-diHpDHA being produced (Table 5 and Supporting Information, Figure 3S). Along with the primary di-HDHA products, minor amounts of tri-HDHA secondary products were produced in the reaction, representing less than 4% of the total and comprising mainly 7,16,17-triHDHA, indicating that the low ratio of 7S,17S-diHDHA to 7S,14S-diHDHA was not due to formation of tri-HDHA products. When 7S-HpDHA was used as the substrate, the products were comparable, with 65% 7S,14S-diHpDHA (i.e. the oxidized form of 7S,14S-diHDHA) and 35% 7S,17S-diHpDHA. This specificity is the opposite of that seen with AA as the substrate for h15-LOX-1, where 90% of the *ω*-6 product, 15-HpETE, is made and only 10% of the *ω*-9 product, 12-HpETE. To confirm the product structures, UV-vis absorbance maxima, C18-reverse phase retention times and MS/MS fragmentations were matched to known patterns for 7S,14S-diHDHA (Supporting Information, Figure 3S), indicating that h15-LOX-1 has a different positional specificity in its reaction with 7S-HDHA than it does with DHA. For comparison, h15-LOX-1 was reacted with DHA to generate 65% of the canonical *ω*-6 product, 17S-HpDHA, with a variety of minor products (22% 14S-HpDHA, 7% 11-HpDHA and 6% 20-HpDHA (Table 5)). The presence of 22% 14S-HpDHA is consistent with oxidation at the *ω*-9 position (10% 12-HpETE from AA), however, the observation of the 10-HpDHA and 20-HpDHA minor products is unusual and indicates that the added length and unsaturation of DHA leads to multiple productive catalytic poses of DHA in h15-LOX-1, similar to that seen with h5-LOX. These rates and product profiles support both Pathways 2 and 3 (Scheme 1) as the most likely in vitro biosynthetic routes for 7S,14S-diHDHA production.

**Table 5.** Distribution of products generated by h15-LOX-1 from DHA, 7S-HDHA and 7S-HpDHA, as determined by LCMS/MS. h15-LOX-1 shows altered positional specificity when reacting with 7S-HDHA and 7S-HpDHA. Note, the abbreviation, “11-product” etc., is used to simplify the complexity of the table labels.

| Substrate | 11-product | 14-product | 17-product | 20-product |
| --- | --- | --- | --- | --- |
| DHA | 7 ±1% | 22 ±5% | 65 ±4% | 6 ±2% |
| 7S-HDHA | 0 | 90 ±6% | 10 ±6% | 0 |
| 7S-HpDHA | 0 | 65 ±8% | 35 ±8% | 0 |

### Reaction of 7S-HDHA and 7S-HpDHA with h15-LOX-2 in *biosynthesis of 7S,17S-diHDHA*

As observed above, h15-LOX-1 only produces a small amount of 7S,17S-diHDHA when reacting with 7S-HDHA. This result raises the possibility that h15-LOX-2 could be the primary LOX isozyme responsible for the generation of 7S,17S-diHDHA (RvD5). In order to study this reaction further, h15-LOX-2 was reacted with DHA, 7S-HDHA and 7S-HpDHA and all three were determined to have comparable rates (Table 6). The *k_cat_* for DHA was 3.0 sec^-1^, the *k_cat_* for 7S-HDHA was 2-fold higher at 5.8 sec^-1^, while the *k_cat_* for 7S-HpDHA was 3.4 sec^-1^. Due to the lower *K_M_* for DHA, the *k_cat_/K_M_* for DHA was 2-fold greater than that of 7S-HDHA (*k_cat_/K_M_* = 0.15 sec^-1^µM^-1^) and 3-fold greater than 7S-HpDHA (*k_cat_/K_M_* = 0.08 sec^-1^µM^-1^). These values indicate that DHA has the fastest rate of substrate capture at low substrate concentrations, *k_cat_/K_M_*, but 7S-HDHA has the fastest rate of product release at high substrate concentrations, *k_cat_* (49). For comparison, the kinetic parameters for h15-LOX-2 and AA are 0.96 sec^-1^ for *k_cat_* and 0.19 sec^-1^µM^-1^ for *k_cat_/K_M_*, similar to published values (15, 43), indicating that at low substrate concentration, 7S-HDHA is a comparable substrate to that of AA, but DHA is preferred. These data support Pathway 5 (Scheme 1) as the most likely *in vitro* biosynthetic route for RvD5 production.

**Table 6.** Steady-state kinetic values for h15-LOX-2 with DHA, 7S-HDHA and 7S-HpDHA.

| Substrate | $k_{cat}$ (sec<sup>-1</sup>) | $k_{cat}/K_M$ (sec<sup>-1</sup>μM<sup>-1</sup>) | $K_M$ (μM) |
| --- | --- | --- | --- |
| AA | 0.96 ± 0.09 | 0.19 ± 0.02 | 5.0 ± 0.3 |
| DHA | 3.0 ± 0.3 | 0.27 ± 0.03 | 11 ± 2 |
| 7S-HDHA | 5.8 ± 0.4 | 0.15 ± 0.02 | 40 ± 4 |
| 7S-HpDHA | 3.4 ± 0.4 | 0.08 ± 0.01 | 41 ± 7 |

To determine the oxylipins produced by h15-LOX-2 from DHA, 7S-HDHA and 7S-HpDHA, the reaction products were isolated and analyzed via LC-MS/MS. The reaction of h15-LOX-2 with DHA produced 95% 17S-HDHA, 3% 14S-HDHA and 2% 20-HDHA (Supporting Information, Figure 4S), similar to its high level of specificity with AA as the substrate (i.e. greater than 95% 15-HpETE). The reaction of 7S-HDHA with h15-LOX-2 produced 100% 7S,17S-diHDHA (Supporting Information, Figure 5S), while reaction with 7S-HpDHA produced 98% 7S,17S-diHDHA and 2% 7S,20-HDHA (Table 7). The 7S,17S-diHDHA produced by h15-LOX-2 from 7S-HDHA was compared with standards and found to have identical UV-vis maxima, LC-MS/MS retention time and fragmentation as 7S,17S-diHDHA (RvD5). It should be noted that no products were observed from the reaction of h15-LOX-2 with either 17S-HDHA or 17S-HpDHA using UV-vis spectroscopy or LC-MS/MS, indicating that 7S,17S-diHDHA is not biosynthesized through the reverse orientation of the substrate. In addition, h15-LOX-2 was not observed to produce the 16,17-epoxide product from 17S-HpDHA, reinforcing its selective reactivity to abstracting only from the *ω*-8 carbon to produce the *ω*-6 oxygenation product.

**Table 7.** Products made from reaction of h15-LOX-2 with DHA-derived oxylipins. DHA, 7S-HDHA, 7S-HpDHA, 17S-HDHA and 17S-HpDHA were reacted with h15-LOX-2, extracted and analyzed via LC-MS/MS. No reaction (no rxn) was detectable after the incubation of h15-LOX-2 with either 17S-HDHA or 17S-HpDHA.

| Substrate | 14-product | 17-product | 20-product |
| --- | --- | --- | --- |
| DHA | 3 ±1% | 95 ±2% | 2 ±1% |
| 7S-HDHA | 0% | 100 ±1% | 0% |
| 7S-HpDHA | 0% | 98 ±1% | 2 ±1% |
| 17S-HDHA | no rxn | no rxn | no rxn |
| 17S-HpDHA | no rxn | no rxn | no rxn |

### Formation of 7S,17S-diHDHA by h5-LOX from 17S-HDHA and 17S-HpDHA

17S-HDHA was reacted with h5-LOX to investigate the rate of 7S,17S-diHDHA synthesis, relative to the production of 7S,17S-diHDHA from h15-LOX-2 and 7S-HDHA. Despite achieving complete turnover of DHA with h5-LOX in ∼1 minute, an identical amount of h5-LOX did not produce an observable reaction with 17S-HDHA or 17S-HpDHA when monitoring at 254 and 270 nm, using UV-vis spectroscopy. However, analyzing the reaction at multiple time points using LC-MS/MS allowed for the estimation of the *V_max_* value. Compared to the reaction of h5-LOX with AA and DHA, the reaction rates with 17S-HDHA and 17S-HpDHA were over 100-fold slower (Table 8). The products of the reaction of h5-LOX with both 17S-HDHA and 17S-HpDHA were approximately 90% 7S,17S-diHDHA, along with minor amounts of 16,17-diHDHA and 10,17-diHDHA. For comparison, similar reactions were performed with h15-LOX-2 and the 7S-oxylipins and the *V_max_* rates were significantly greater than observed with h5-LOX (Table 8), indicating that Pathway 6, Scheme 1 is not efficient in vitro. As with h15-LOX-2, it is possible that 7S,17S-diHDHA could also be biosynthesized through a reverse binding orientation to h5-LOX, however, no reaction was observable between h5-LOX and 7S-HDHA or 7S-HpDHA, using UV-vis spectroscopy. When analyzed by LC-MS/MS, 7S,17S-diHDHA was observed, but the reaction rates of h5-LOX with 7S-HDHA and 7S-HpDHA were over 1000-fold slower than with AA and DHA and were considered inconsequential (i.e. no reaction).

**Table 8.**
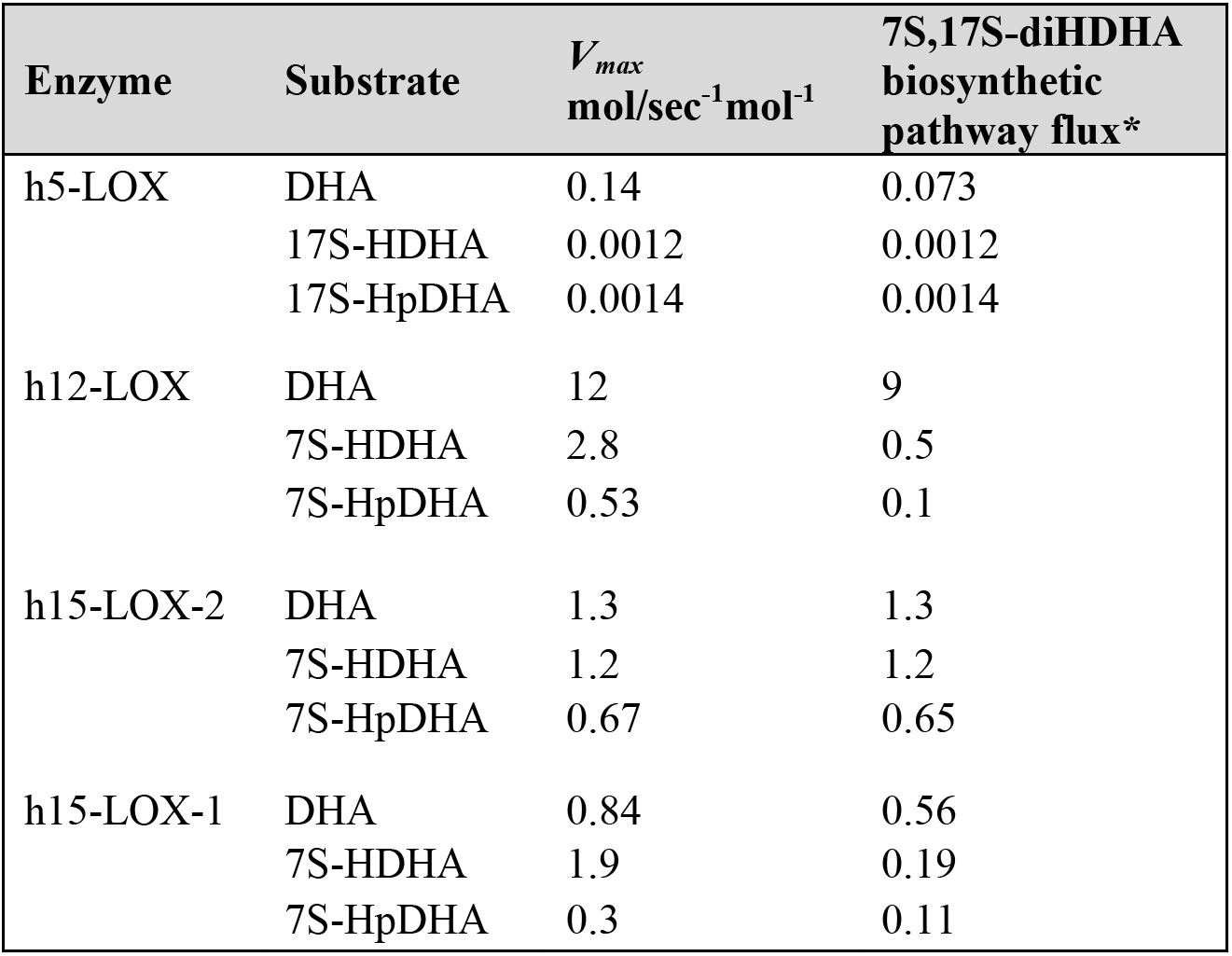
V_max_ values of reaction steps in the biosynthesis of RvD5 were calculated at 10 µM substrate concentration for comparison. *Biosynthetic flux is calculated by multiplying each *V_max._* by the percentage of total product from that reaction that serves as substrate for the next step in the synthesis of 7S,17S-diHDHA. Reactions with DHA represent the first of two biosynthetic steps, while reactions with oxylipins represent the second.

### Chiral chromatography characterization of 7S,14S-diHDHA and 7S,17S-diHDHA (RvD5)

We predicted that the oxygen on C14 of 7S,14S-diHDHA is in the S-configuration because h15-LOX-1 typically performs stereospecific oxygenation generating the S-configured product when the substrate enters the active site methyl-end first. To determine the relative stereochemistry of the 7S,14S-diHDHA produced by reacting 7S-HDHA with h15-LOX-1, the products of this reaction were isolated, reduced and compared to standards via reverse phase chiral chromatography. MaR1 and 7-epi*-*MaR1 standards (Cayman Chemicals) and the 7S,14S-diHDHA produced by h15-LOX-1 from 7S-HDHA eluted at distinct times when analyzed via chiral HPLC (Figure 2). 7-epi-MaR1 eluted with a RT of 45.5 min, MaR1 with a RT of 37.9 min and the enzymatically-synthesized 7S,14S-diHDHA eluted with a RT of 22.4 min. All three compounds shared identical fragmentations when analyzed via LC/MS/MS. As no 7R,14S-EZE-diHDHA or 7S,14S-EZE-diHDHA standards are commercially available, resolution of these two isomers from one another was confirmed by reacting h15-LOX-1 with a racemic mixture of 7R/S-HDHA, resulting in two peaks at 20.8 and 22.4 min with identical fragmentations and UV-spectra. Since h5-LOX also typically generates product in the S-configuration from substrates that enter the active site methyl-end first, the reaction of h5-LOX with 14S-HDHA is predicted to produce 7S,14S-diHDHA. A comparison of this product with that generated by h15-LOX-1 from 7S-HDHA revealed a chiral retention time of 22.4 min, identical to that of the 7,14-diHDHA produced by h15-LOX-1 from 7S-HDHA. This indicates that both oxylipins are 7S,14S-diHDHA, just produced through a different order of LOX biosynthetic steps. While it is conceivable that both h15-LOX-1 and h5-LOX change their product profile and generate the R-configuration oxylipin, this is not the case based on the chiral column results. It should be noted that a small 7R,14S peak is generated by the reaction of 14S-HDHA with h5-LOX. In addition, a small 7R,14S-diHDHA peak is generated by the reaction of 7S-HDHA with h15-LOX-1, indicating that the 7S-HDHA synthesized by h5-LOX from DHA contained a small contaminate of 7R-HDHA. Together, these two observations indicate that while the major product of h5-LOX is the S-configured product, a small amount of the R-product is also made.

**Figure 2.**
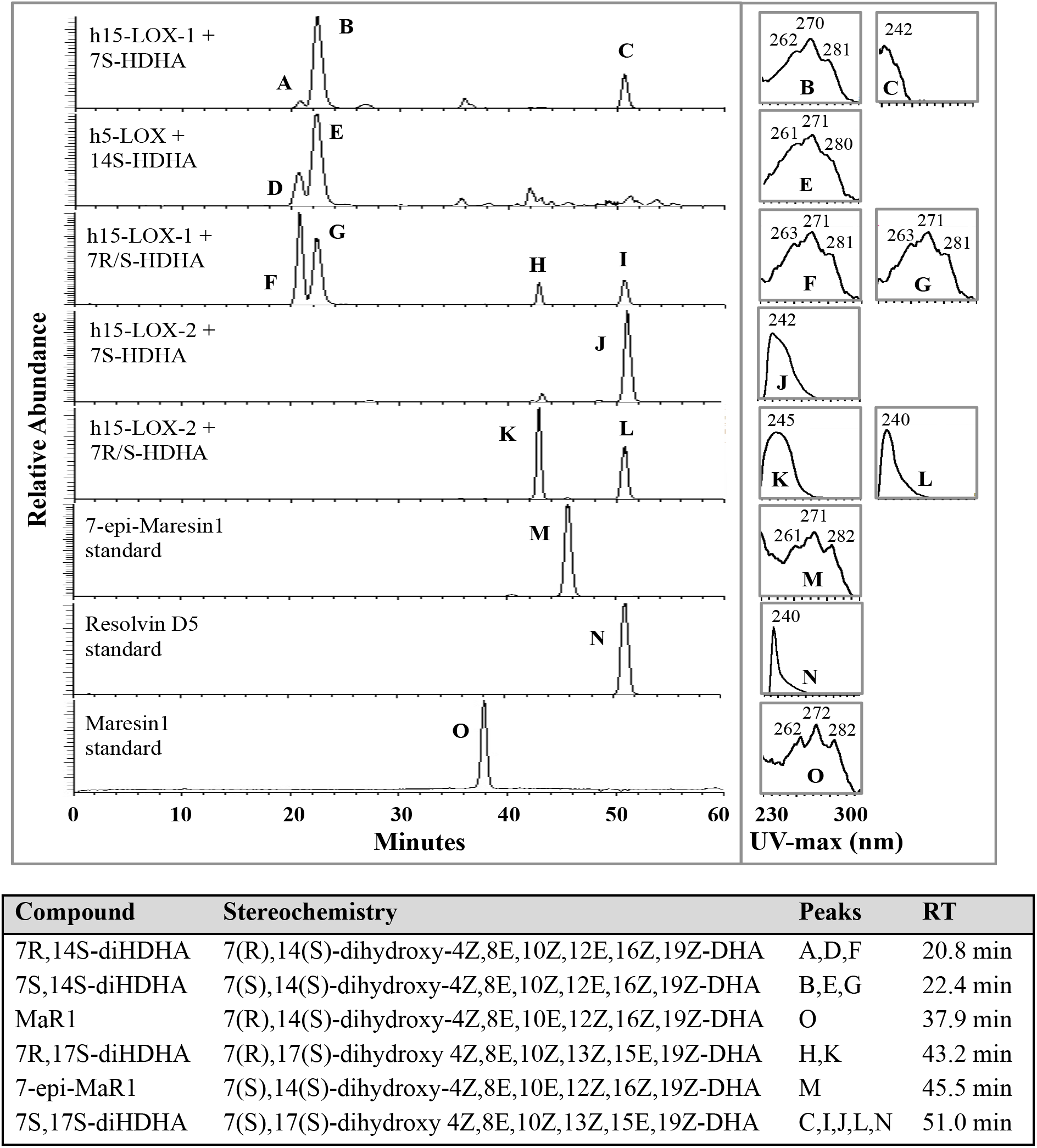
Chiral chromatograms and UV-maxima of 7,14-diHDHA and 7,17-diHDHA isomers. The products formed by h15-LOX-1 and h15-LOX-2 from 7S-HDHA and 7R/S-HDHA and by h5-LOX from 14S-HDHA were analyzed via Chiral LC-MS/MS and UV spectra and compared to MaR1, 7-epi-MaR1 and RvD5 standards. All 7,14-diHDHA isomers contained a central peak at ∼270 nm, surrounded by shoulders at ∼260 nm and ∼280 nm, consistent with the presence of a conjugated triene. Shoulders of equal intensity at 281 nm and 261 nm are indicative of an EEZ configuration, while a more intense shoulder at 260 nm compared to 280 nm indicates the EZE configuration. 7,17-diHDHA isomers contain two conjugated dienes, with UV-maxima of 240-245 nm, distinguishing it from 7,14-diHDHA.

Along with chiral retention times, UV spectra were compared to further differentiate the double bond geometry of the various products. All 7,14-diHDHA products contained a central peak at ∼270 nm, surrounded by shoulders at ∼260 nm and ∼280 nm, consistent with the presence of a conjugated triene. In Mar1 and 7-epi-MaR1, the shoulders at 281 nm and 261 nm were of equal intensity, indicative of a conjugated triene with EEZ configuration (50–52). In contrast, the 7,14-HDHA produced enzymatically showed a more intense shoulder at 260 nm than at 280 nm, indicating the EZE configuration (53). It should be noted that we have attempted to determine the absolute stereochemistry of 7S,14S-diHpDHA by generating the double Mosher derivative, but unfortunately low yields from enzymatic synthesis and NMR peak overlap prohibits the assignment of the molecules’ stereochemistry.

To determine the relative stereochemistry of the 7,17-oxylipin produced by h15-LOX-2 and 7S-HDHA, the molecule was compared to an RvD5 standard using reverse-phase, chiral HPLC coupled to MS/MS. The chiral HPLC retention time, MS/MS fragmentation and UV-spectra of the 7,17-oxylipin produced by h15-LOX-2 from 7S-HDHA matched those of the RvD5 standard (Cayman Chemicals) (Figure 2), strongly suggesting its identity to be 7S,17S-diHDHA (RvD5). 7S,17S-diHDHA and 7R,17S-diHDHA were resolved under these conditions and demonstrated UV-maxima of 240-245 nm, indicative of two separate conjugated-dienes

### Molecular Modeling of DHA and 7S-HDHA bound to h15-LOX-1

The shift in product profile seen between h15-LOX-1 reacting with DHA and 7S-HDHA suggests that 7S-HDHA has a different binding mode than DHA, and therefore molecular modeling was employed to assess this possibility. Extra-precision Glide scores were used to approximate ligand binding free energy, with lower negative scores representing tighter binding (54). Figure 3 shows the InducedFit docking poses of DHA and 7S-HDHA with h15-LOX-1, with corresponding extra-precision docking scores for these poses of -9.5 and -10.8, respectively. In the docking model, the carboxylate groups of both DHA and 7S-HDHA form hydrogen bonds with the side chain of Arg402 and the hydrophobic tails of both substrates are buried deep in the hydrophobic pocket created by residues Phe352, Ile417 and Ile592. However, a key difference is that the C7 hydroxyl group of 7S-HDHA forms a hydrogen bond with the backbone carbonyl oxygen of Ile399. This difference in binding between DHA and 7S-HDHA is manifested in their distances between the iron-hydroxide oxygen atom and the hydrogen on the reactive carbons, C12 and C15. The modeling data indicate that for 7S-HDHA, the C12 pro-S hydrogen is markedly closer (2.6 Ang) to the iron-hydroxide moiety than C15 hydrogen (5.9 Ang) (Table 9). Considering that C12 hydrogen atom abstraction leads to 7S,14S-diHDHA, while C15 hydrogen atom abstraction leads to 7S,17S-diHDHA, these docking results are consistent with the enzymatic results. For DHA, the distance for the C15 pro-S hydrogen (3.8 Ang) is slightly shorter than that of C12 (4.1 Ang), consistent with the experimental results, but the distance difference is not as distinct as that for 7S-HDHA, possibly due to the more homogeneous hydrophobic nature of DHA compared to 7S-HDHA. Interestingly, there is no difference between the pro-S and pro-R hydrogens indicating the limitation of the docking model.

**Figure 3.**
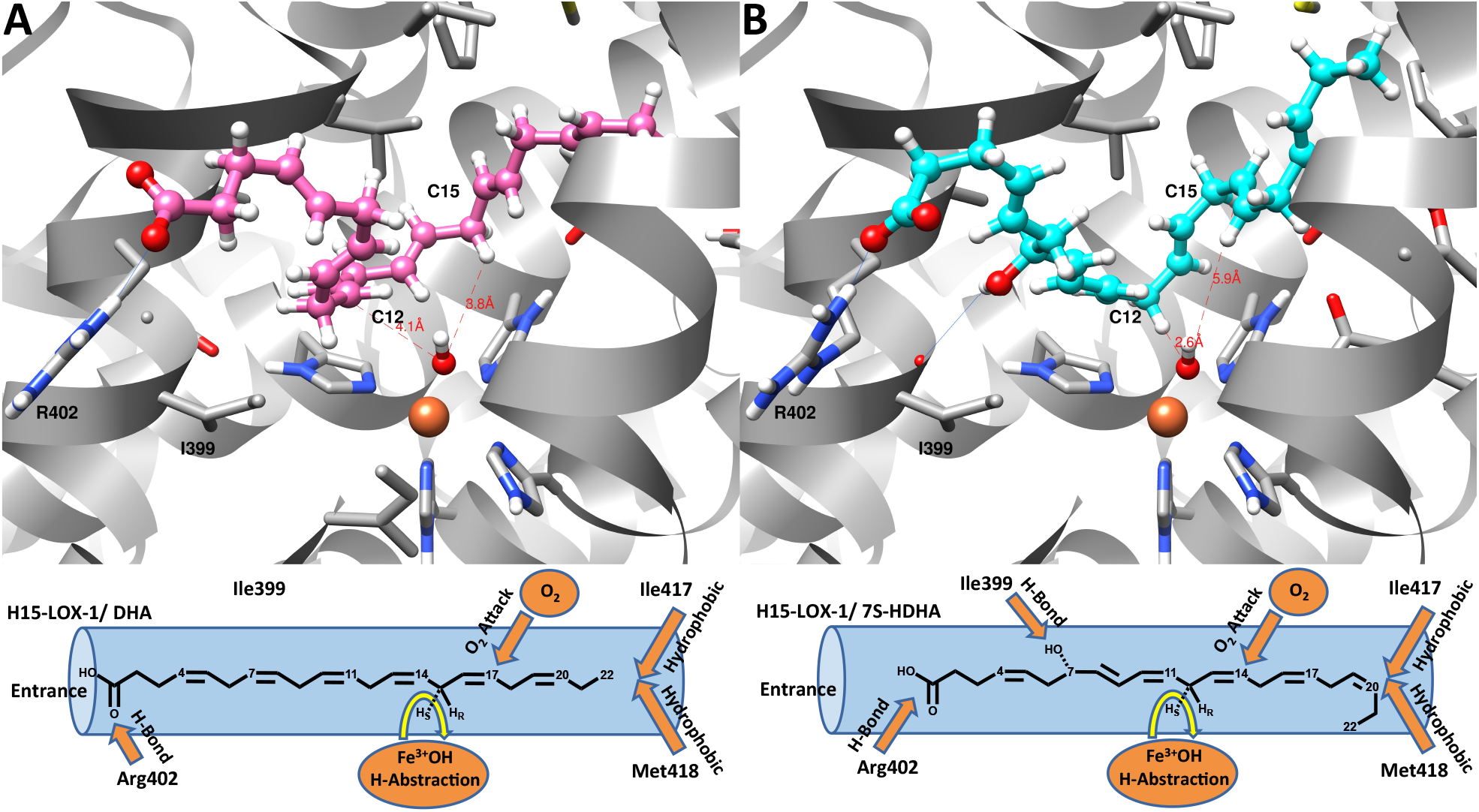
InducedFit docking poses of (A) DHA and (B) 7S-HDHA against the active site of h15-LOX1. DHA and 7S-HDHA atoms, hydroxide ion and metal ion are shown in ball-and-stick representation. Protein residues are shown in stick representations. Carbon atoms of DHA, 7S-HDHA and protein are shown in pink, cyan and gray, respectively; nitrogen, oxygen and hydrogen atoms are shown respectively in blue, red and white. Fe^3+^ is shown in orange. Hydroxide oxygen to C12-hydrogen and C15-hydrogen distances of DHA and 7S-HDHA are also shown. Below each docking structure, a cartoon representation is presented for positional emphasis.

**Table 9.** Distances between hydroxide-ion oxygen atom and the pro-S hydrogens of the reactive carbons, C12 and C15, of DHA and 7S-HDHA against h15-LOX-1.

| **Substrate** | **Hydroxide**<br>**oxygen to pro-S**<br>**C12-hydrogen**<br>**distance (Å)** | **Hydroxide**<br>**oxygen to pro-S**<br>**C15-hydrogen**<br>**distance (Å)** |
| --- | --- | --- |
| **DHA** | 4.1 | 3.8 |
| **7S-HDHA** | 2.6 | 5.9 |

### Molecular modeling of DHA and 7S-HDHA bound to h15-LOX-2

For h15-LOX-2, we chose the extra-precision (XP) scoring function for Glide docking within the InducedFit docking module, which allowed for extensive sampling of the substrate conformations during docking. The XP docking score of the low-energy docking pose of DHA and 7S-HDHA against h15-LOX-2 are -7.09 and -9.31 respectively and their docking poses are shown in Figure 4. Both substrates bind into the active site tail first, utilizing a U-shaped binding mode. The carboxylate groups of both DHA and 7S-HDHA make a salt-bridge interaction with the R429 residue on the helix *α*12. In contrast, h15-LOX-1 utilizes R402 (helix *α*11) to hydrogen bond with the carboxylate group of the substrate, possibly leading to their different product specificities. The hydrophobic tails of both substrates are buried deeply in the hydrophobic pocket created by residues F365, L420, I421, V427, F438, and L607. The hydroxyl group of 7S-HDHA makes a hydrogen-bond to the backbone carbonyl oxygen of L419. The distance between reactive pro-S hydrogen of C15 and the oxygen atom of the hydroxide ion is 4.1Å and 2.7Å for DHA and 7S-HDHA, respectively (Table 10). These distances are shorter than the pro-S hydrogen of C12 and the oxygen atom of the hydroxide ion for DHA and 7S-HDHA (7.2 Å and 4.0 Å, respectively) (Table 10). These two binding modes are consistent with the enzymatic reaction involving abstraction of the pro-S hydrogen from C15 for both DHA and 7S-HDHA, to produce the 17-product exclusively (Table 10) (55–59) and not the 14-product from C12 hydrogen abstraction.

**Figure 4.**
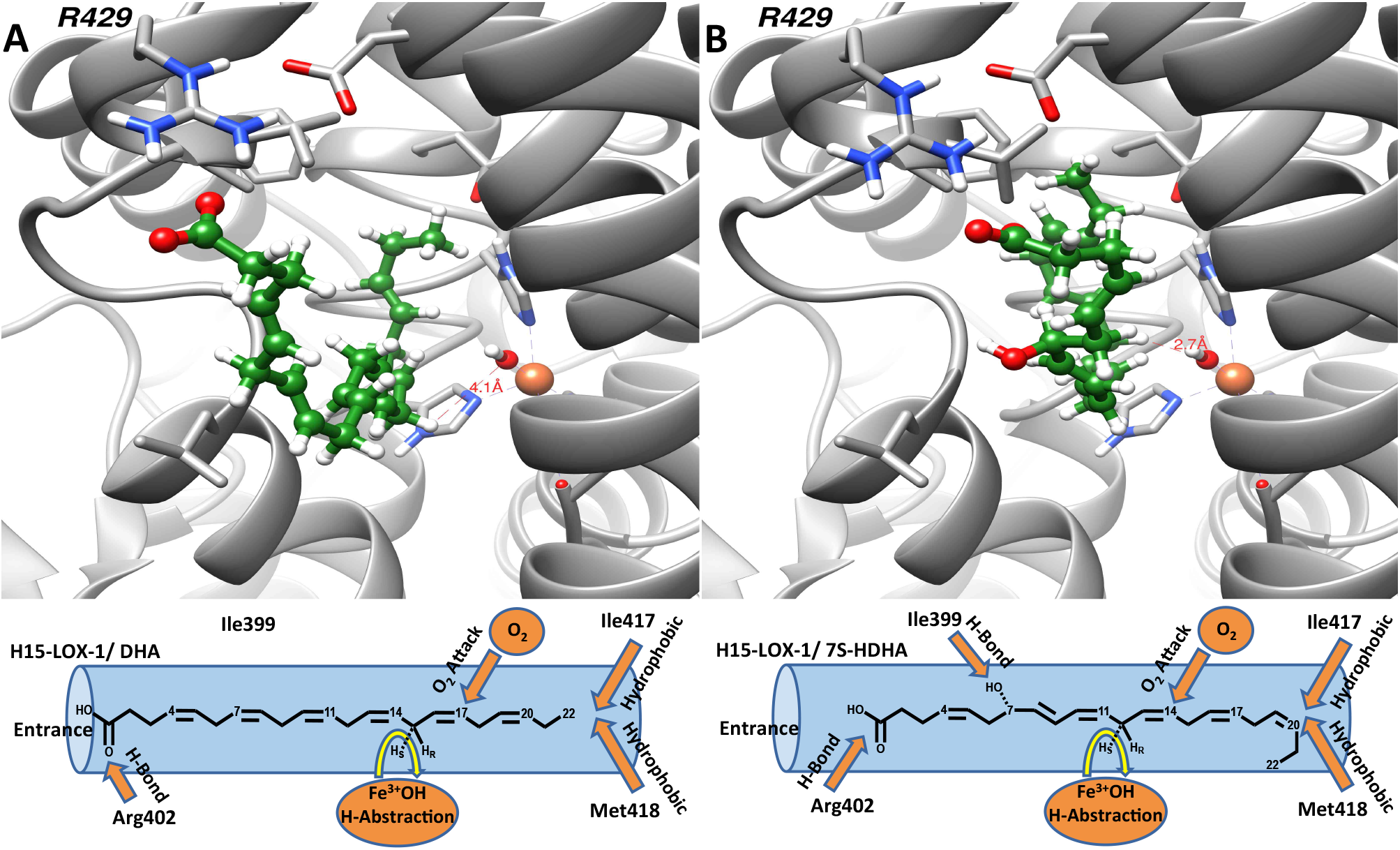
InducedFit docking poses of (A) DHA and (B) 7S-HDHA against the h15-LOX-2 active site. DHA and 7S-HDHA atoms, hydroxide ion and metal ion are shown in ball-and-stick representation and protein residues are shown in stick representation. Carbon atoms of DHA and 7S-HDHA are shown in green color and protein are shown in gray color; nitrogen, oxygen and hydrogen atoms are shown respectively in blue, red and white colors. Fe^3+^ ion is shown in orange. The distance between reactive pro-S hydrogen atom of C15 and the oxygen atom of the hydroxide ion is 4.1Å and 2.7Å for DHA and 7S-HDHA respectively

**Table 10.** Distances between hydroxide-ion oxygen atom and the pro-S and pro-R hydrogens of the reactive carbons, C12 and C15, of DHA and 7S-HDHA with h15-LOX-2.

| Substrate | Hydroxide oxygen to pro-S C12-hydrogen distance (Å) | Hydroxide oxygen to pro-S C15- hydrogen distance (Å) |
| --- | --- | --- |
| DHA | 7.2 | 4.1 |
| 7S-HDHA | 4.0 | 2.7 |

### Effect of 7S,14S-diHDHA and RvD5 on Platelets

Previous studies have shown that DHA and the h12-LOX-derived oxylipins of DHA have antiplatelet effects (60). Therefore, to determine the effect of RvD5, 7S,14S-diHDHA and related maresin isomers on platelet activation, washed platelets were treated with oxylipins in half-log increments and then stimulated with collagen (0.25 µg/mL) in an aggregometer. 7S,14S-diHDHA fully inhibited platelet aggregation at 10 µM, while reducing aggregation by 65% at 3 µM. (Figure 5). Platelets treated with 3 and 10 µM of RvD5 had an approximate reduction in aggregation of 65% and 90%, respectively, while MaR1 and 7-epi-MaR1 were the least potent, reducing aggregation by approximately 70% at 10 µM. 12S-HETrE, a h12-LOX-derived oxylipin of DGLA with known antiplatelet activity (61–63), displayed slightly greater potency than 7S,14S-diHDHA and 7S,17S-diHDHA (RvD5), reducing collagen-mediated platelet aggregation by 68% at 1 µM. Together this data demonstrates that 12-HETrE was more potent than both 7S,17S-diHDHA (RvD5) and 7S,14S-diHDHA for inhibiting collagen-induced platelet aggregation, however at 10 µM, all three molecules inhibited aggregation completely. It should be noted that these conditions (0.25 µg/mL collagen as activator) are more sensitive to inhibition than our previously published conditions (20 µM PAR1-AP as agonist) (62) and therefore demonstrate a greater effect on aggregation at lower concentrations of 12S-HETrE.

**Figure 5.**
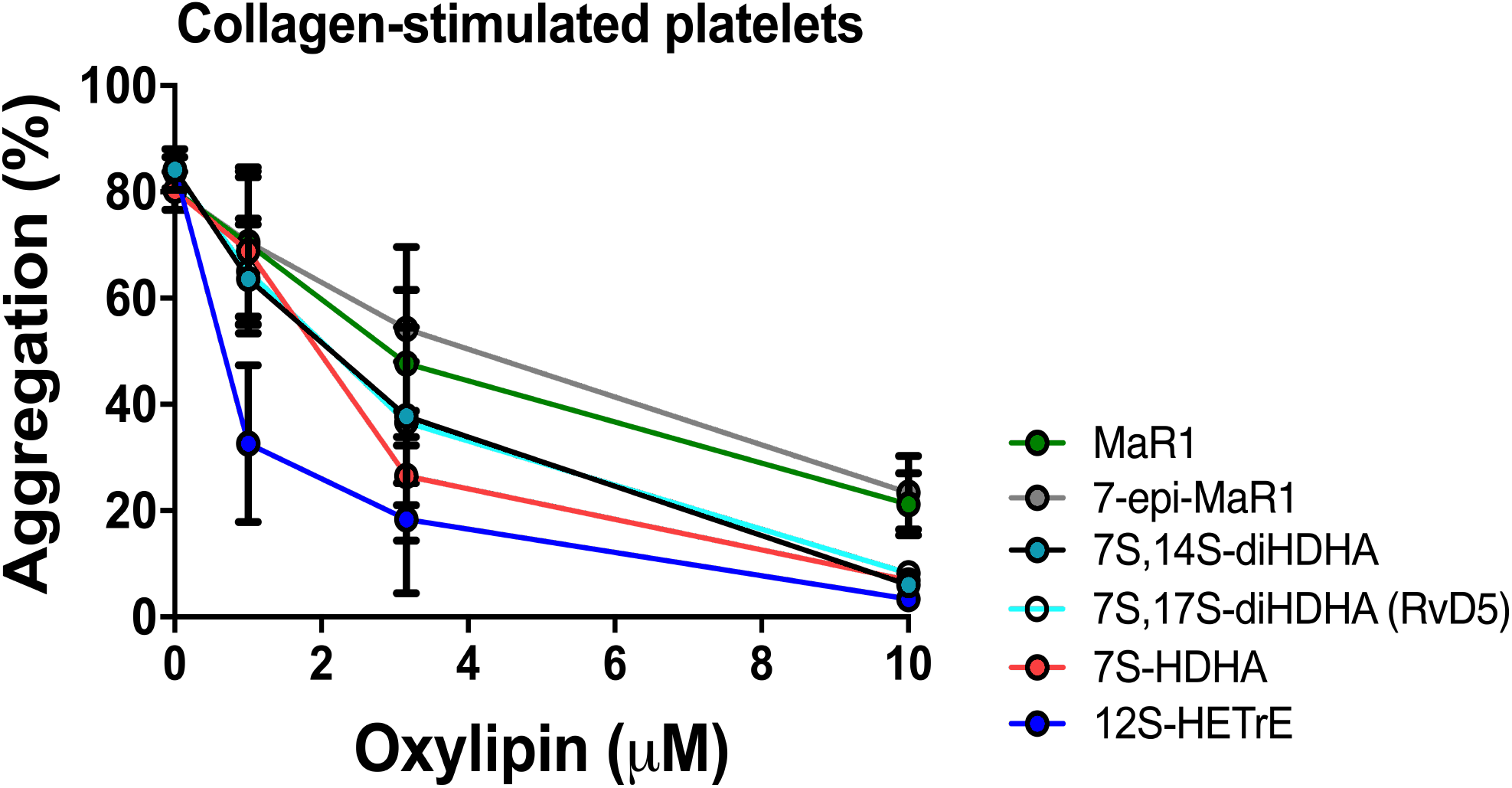
The effect of MaR1, 7-epi-MaR1, 7S,14S-diHDHA, 7S,17S-diHDHA (RvD5), 7S-HDHA, 12S-HETrE, Mar1 and 7-epi-Mar1 on aggregation of activated platelets. Washed human platelets were treated with vehicle (DMSO) or MaR1, 7-epi-MaR1, 7S,14S-diHDHA, 7S,17S-diHDHA, 7S-diHDHA, or 12S-HETrE in half-log increments ranging from 1-10 µM for ten minutes and then stimulated with collagen (0.25 µg/mL) in an aggregometer. Note, the data for 7S,14S-diHDHA and 7S,17S-diHDHA are nearly identical and thus overlap significantly. Data represent the mean ± S.E.M of 5 independent experiments. Statistical analysis was performed using one-way ANOVA with Dunnett’s test comparing the aggregation of platelets treated with each concentration of oxylipin to the aggregation of vehicle-treated platelets. **P<0.01, ***P<0.001

### Platelet Lipidomics with 7S-HDHA

The reduction in aggregation seen in platelets incubated with 7S-HDHA could be caused by either the 7S-HDHA itself or by a biosynthetic product produced by platelets during incubation. To investigate the latter possibility, platelets were incubated with 7S-HDHA and analyzed for the presence of 7S-HDHA metabolites using LC-MS/MS. Although large amounts of unreacted 7S-HDHA were detectable in the samples, no levels of any di-HDHA’s or tri-HDHA’s were detectable. As a control, platelets incubated with 5S-HETE were also analyzed for the presence of 5S-HETE metabolites. Although large levels of 12-HETE were detectable from endogenous AA, no appreciable levels of 5S-HETE/ h12-LOX metabolites were detectable down to 2 ng/mL. This is an unexpected result since h12-LOX is capable of reacting with 7S-HDHA *in vitro* to produce 7S,14S-diHDHA. The fact that no 7S,14S-diHDHA is produced indicates both that 7S-HDHA is most likely the bioactive species and that 7S-HDHA does not react appreciably with h12-LOX in the cellular milieu of the platelet.

## DISCUSSION

### Biosynthesis of 7S,14S-diHDHA

7S,14S-diHDHA is an analogue of MaR1 (17) and is proposed to have a distinct bio-synthetic pathway from MaR1. Its structure suggests three possible routes for its biosynthesis. Pathway 1 involves that oxidation of DHA at C14 by h12-LOX, followed by oxidation at C7 by h5-LOX, however, the *in vitro* data of this work suggests this is a very unfavorable biosynthetic route. Although DHA is a good substrate for h12-LOX, forming 14S-HpDHA at a rate similar to 12S-HpETE formation from AA (44), the second step in this pathway is kinetically unfavorable. Compared with DHA, the *V_max_* of h5-LOX with 14S-HDHA is ∼368 fold slower (Table 2).

For Pathway 2 in Scheme 1, DHA is oxidized at C7 by h5-LOX followed by oxidation at C14 by h12-LOX. In comparison to Pathway 1, Pathway 2 is a kinetically favorable in-vitro biosynthetic route. The first step, formation of 7S-HpDHA from DHA by h5-LOX, occurs rapidly with a *V_max_* comparable to that of AA kinetics. The second step is also favorable, as the reaction of h12-LOX with 7S-HDHA has *k_cat_* and *k_cat_*/*K_M_* values that are within 10-fold the rate of DHA. The slowest step in this pathway (i.e the rate determining step) is over 350 times faster than the reaction of h5-LOX with 14S-HDHA.

Interestingly, Pathway 2 also presents a possible novel pathway to biosynthesizing 7S,17S-diHDHA (RvD5) with h12-LOX. The production of 7S,17S-diHpDHA is 18% of that of 7S,14S-diHpDHA, which is a significant percentage given the potency of RvD5 (25). The result also indicates a product promiscuity rarely seen for h12-LOX. h12-LOX produces over 95% 12S-HpETE from AA, so to produce 18% 7S,17S-diHpDHA is noteworthy. Another interesting aspect of this result is the mechanism of 7S,17S-diHpDHA formation. For 7S,17S-diHpDHA to form, h12-LOX must abstract from C15 of 7S-HpDHA, which indicates that the methyl tail of 7S-HpDHA is not positioned as deep in the active site pocket as that for 7S,14S-diHpDHA formation. These data suggest that the alcohol on C7 of 7S-HpDHA could interact with the active site and prevent the full penetration of the methyl tail into the bottom of the active site pocket.

The final biosynthetic route examined, Pathway 3, is the oxidation of DHA at C7 by h5-LOX, followed by oxidation at C14 by h15-LOX-1, and is a non-canonical pathway. h15-LOX-1 reacts with DHA and produces a majority of 17S-HpDHA (65%), with a smaller amount of 14S-HpDHA (22%). As shown with AA (64), product specificity is dictated by the methyl end of the fatty acid, with the majority of oxygenation occurring on the *ω*-6 carbon (C15 for AA and C17 for DHA), with a minority occurring on the *ω*-9 carbon (C12 for AA and C14 for DHA). Therefore, it is reasonable to expect a similar distribution of products with 7S-HDHA as the substrate, however this is not the case. Pathway 3 produces over 9–fold more of 7S,14S-diHDHA than 7S,17S-diHDHA. In addition, this unusual product pathway is kinetically favorable *in vitro*. Both the first step, formation of 7S-HpDHA from DHA by h5-LOX, and the second, reaction of 7S-HDHA with h15-LOX-1 occur relatively rapidly. The *k_cat_* for h15-LOX-1 and 7S-HDHA is 3.2-fold faster than that of DHA and the *k_cat_/K_M_* for DHA and 7S-HDHA are nearly identical. These results indicate that, although the hydroxyl group on C7 has a large effect on the positional specificity of the enzyme, it has little effect on catalysis, leading to a favorable overall kinetic condition. In summary, if we consider the product of both the rate and the percent product formation (i.e. the biosynthetic flux, Scheme 2), the data indicates that the most efficient pathway for making 7S,14S-diHDHA is thru h5-LOX and then h12-LOX or h15-LOX-1 and not vice versa since the reaction of h5-LOX with 14-HpDHA is markedly slower and hence rate-limiting.

**Scheme 2.**
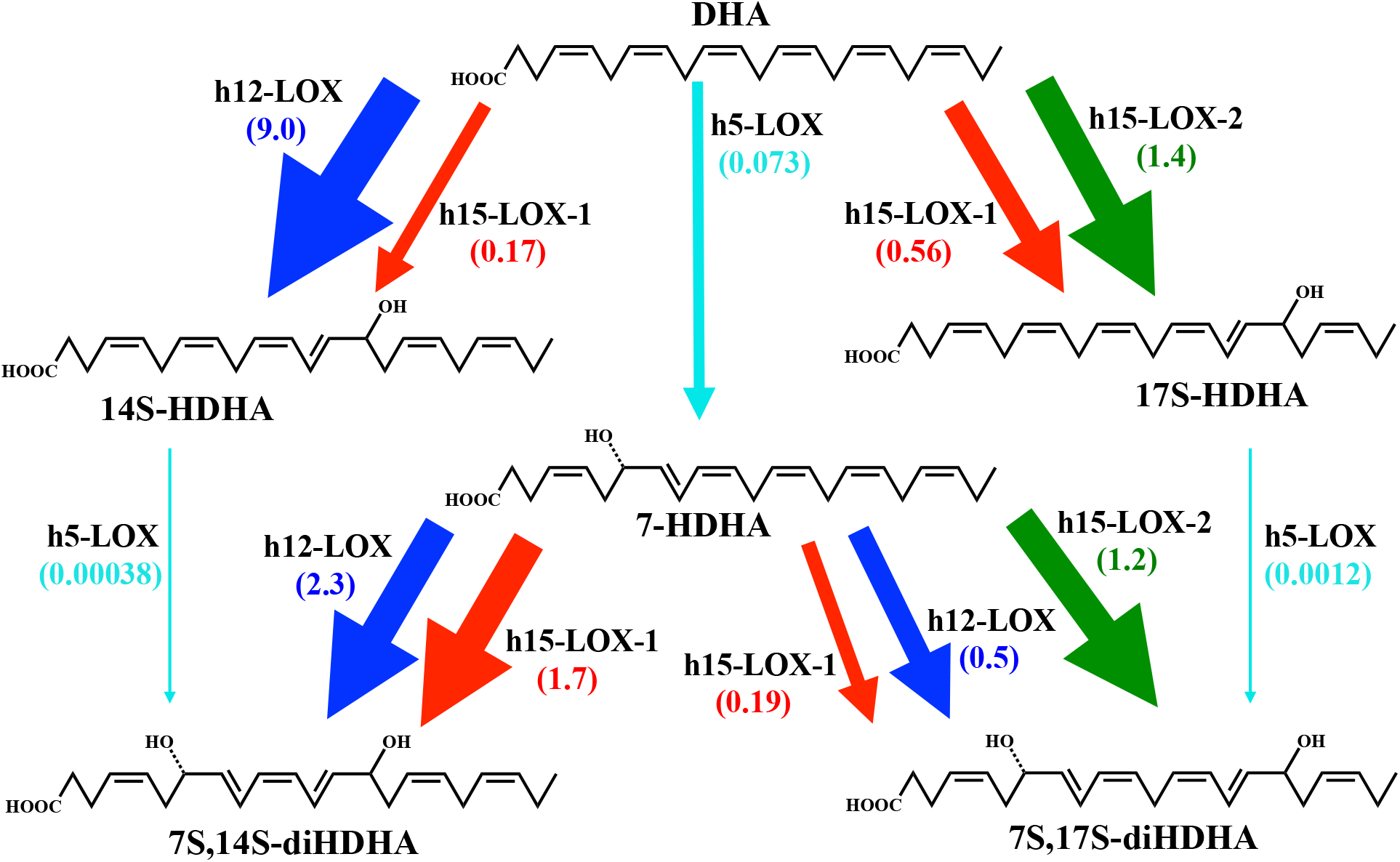
Comparison of pathways for the biosynthesis of RvD5.and 7S,14S-diHDHA. The total biosynthetic flux for the reactions of h5-LOX, h12-LOX and h15-LOX-1 with DHA, 14S-HpDHA and 7S-HpDHA are compared by multiplying *V_max_* values (mol/sec^-1^mol^-1^) at 10 nM substrate by the percent product for each reaction. In all pathways, turnover by h5-LOX is the rate-limiting step and we have assumed that the hydroperoxides are reduced to the alcohols.

Although changes in the positional specificity of LOXs have been demonstrated before (19), the finding that h15-LOX-1 primarily oxygenates C17 on DHA, but C14 on 7S-HDHA is surprising. In order to obtain a better understanding of this binding event, DHA and 7S-HDHA were docked to a model of h15-LOX-1. In the case of DHA, both C12 and C15 are at a reasonable distance for hydrogen abstraction (65), whereas in case of 7S-HDHA, C12 is closer to the metal ion than C15, supporting the experimental results (Table 9). The model explains this result by having the hydroxyl group at C7 in 7S-HDHA forming a hydrogen bond with the backbone carbonyl of I399, which subsequently positions C12 closer to the metal ion. Without a similar restraint, the flexibility of DHA positions both C12 and C15 close to the metal ion, helping to explain the greater number of products made by h15-LOX-1 from DHA compared with AA. This model suggests that 7S-HDHA is positioned further inside the active site than DHA, overcoming steric interactions to position C12 for hydrogen atom abstraction (Figure 3). What is remarkable is that this is the opposite from what appears to occur with 7S-HDHA in h12-LOX. In h12-LOX, the C7 alcohol holds the substrate back from the bottom of the pocket to abstract a hydrogen atom closer to the methyl tail generating the 7S,17S-diHDHA, while in h15-LOX-1, the C7 alcohol positions the substrate deeper into the active site generating 7S,14S-diHDHA. We are currently investigating the active sites of both h12-LOX and h15-LOX-1 with mutagenesis to better understand the binding requirements and differences between h12-LOX and h15-LOX-1 in more detail.

### Biosynthesis of 7S,17S-diHDHA (RvD5)

The discovery of Pathway 3 as a relevant route to the biosynthesis of 7S,14S-diHDHA raises questions with respect to the biosynthetic route to making 7S,17S-diDHA (RvD5). Previously, 7S,17S-diDHA (RvD5) was proposed to be biosynthesized from DHA in two steps; initial oxygenation of C7 by h5-LOX, followed by oxygenation of C17 by h15-LOX-1 (24, 26). This hypothesis was based on the prevalence of h15-LOX-1 in the inflammasome and the preference of h15-LOX-1 to oxygenate DHA at C17. However, our work shows that h15-LOX-1 shifts its positional specificity when reacting with 7S-HDHA *in vitro*, mainly creating 7S,14S-diHDHA and very little of 7S,17S-diHDHA. This marked change in substrate specificity indicates that another LOX, such as h15-LOX-2, could be responsible for the biosynthesis of RvD5.

h15-LOX-2 reacts favorably with DHA, manifesting a *k_cat_* of 3.0 sec^-1^, which is comparable to values published for AA EPA, DGLA and GLA (66), indicating h15-LOX-2 maintains similar rates of product release (*k_cat_*) across a wide range of fatty acid substrates (49). Since the reaction of h15-LOX-2 with DHA produces 95% 17S-HDHA, and the fact that h15-LOX-1 specific inhibitors do not block production of 17S-HDHA in neutrophils and eosinophils (41), it is possible that the likely source of 7S,17S-diHDHA is h15-LOX-2. In fact, 7S-HDHA and 7S-HpDHA are viable substrates of h15-LOX-2, with greater *k_cat_* values but lower *k_cat_*/*K_M_* values than DHA, making them less reactive substrates than DHA at low concentration with h15-LOX-2. Previous work comparing the reaction of AA and 5S-HpETE with h15-LOX-2 demonstrated a similar trend, with 5S-HpETE having a greater *k_cat_* but a smaller *k_cat_*/*K_M_* than AA (15). Considering that hydrogen atom abstraction is generally the first irreversible rate-determining step for *k_cat_* and *k_cat_*/*K_M_* for fatty acid substrates, these results suggest that the rate of oxylipin binding is markedly slower relative to fatty acid binding for h15-LOX-2 (15).

With respect to the biosynthetic pathway of RvD5, the fact that RvD5 contains two hydroxyls on C7 and C17 suggests three likely routes for the biosynthesis of RvD5 from DHA (Pathways 4, 5 and 6, Scheme 1). The first route could be the oxygenation at C7 by h5-LOX followed by oxygenation of C17 by a 15-LOX isozyme (Pathway 4 or 5), either h15LOX-1 or h15-LOX-2. The second route is the reverse, with oxygenation of C17 by a 15-LOX isozyme, followed by oxygenation of C7 by h5-LOX (Pathway 6). In the current study, we find that h5-LOX reacts well with DHA, however, it reacts significantly slower with 7S-HDHA and 17S-HDHA. For a direct comparison, the *V_max_* at 10 µM substrate was determined to be 0.14 mol/sec^-1^mol^-1^ for h5-LOX with DHA (Table 8), which is over 100 times greater than the rate observed with 17S-HDHA (*V_max_* = 0.0012 mol/sec^-1^mol^-1^ at 10 µM). This is consistent with the low reactivity of h5-LOX and 14S-HDHA, indicating that h5-LOX does not react well with oxygenated derivatives of DHA and brings into question Pathway 6 as a viable *in viv*o biosynthetic route. Pathway 4 also appears unfavorable since the current work indicates that h15-LOX-1 reacts with 7S-HDHA to produce 90% 7S,14S-diHDHA (*vide supra*). It therefore appears that h15-LOX-1 may not contribute much to the *in vitro* biosynthesis of RvD5. h15-LOX-2, however, reacts well with 7S-HDHA (*V_max_* = 1.2 mol/sec^-1^mol^-1^ at 10 µM) (Table 8), indicating that the rate of oxygenation at C7 by h5-LOX followed by oxygenation of C17 by h15-LOX-2 is a markedly faster pathway than the rate of oxygenation of C17 by h15-LOX-2 followed by oxygenation of C7 by h5-LOX. Therefore, if we calculate the biosynthetic flux (i.e. the rates multiplied by the percent product formation (Scheme 2)), it is apparent that the preferred pathway of RvD5 production *in vitro* is thru h5-LOX and then h15-LOX-2.

Previously, it was shown that h15-LOX-2 oxygenated 5S-HpETE, but not 12S-HpETE or 15S-HpETE (15). Together with the current work, this suggests that the h15-LOX-2 active site tolerates substrates which are oxygenated close to the carboxylate end, but not those substrates oxygenated closer to the methyl end. This is consistent with the substrate-binding model, where the methyl-end of the substrate binds at the bottom of the active site (64, 67, 68), allowing for the abstraction of a hydrogen atom from C15 and the subsequent oxidation at C17. In order to understand this binding event in more detail, DHA and 7S-HDHA were modeled into the active site of h15-LOX-2. Both substrates bind in a tail first, U-shaped binding mode with their carboxylate groups forming a salt-bridge with R429 on the helix *α*12. Their hydrophobic tails are buried deep in a hydrophobic pocket created by residues F365, L420, I421, V427, F438 and L607 (Figure 4), and the reactive pro-S hydrogen of C15 of 7S-HDHA is 2.7Å from the active site hydroxide-ferric moiety. This binding mode is consistent with the abstraction of the pro-S hydrogen atom from C15 to produce 7S,17S-diHpDHA. Parenthetically, the pro-S hydrogen is also abstracted in the formation of 12-HETE, 5-HETE, and 15-HETE by h12-LOX, h5LOX and h15-LOX-1 respectively (55–59).

### Biological Consequences

If we consider that the *in vitro* biosynthetic pathways outlined above for 7S,14S-diHDHA and 7S,17S-diHDHA could be viable in *in vivo* systems and the fact that both molecules are biologically active, then a few cellular consequences should be discussed. First, the ability of h15-LOX-1 to produce 7S,14S-diHDHA from 7S-HDHA opens up the possibility for involvement of new cells types not previously considered in 7S,14S-diHDHA formation. For example, cellular interactions within blood clots are proposed to lead to the synthesis of SPM through the process of transcellular biosynthesis (27, 28), therefore, cell types that contain h5-LOX and h15-LOX-1 may participate in the transcellular biosynthesis of 7S,14S-diHDHA. Also, h15-LOX-1 is expressed at high levels in macrophages, but it is not found in blood vessel endothelium or in platelets (69–71), while h5-LOX is expressed at high levels in neutrophils (72–74). As macrophages and neutrophils are often co-localized at sites of inflammation and thrombosis, these two cell types may be involved in the transcellular biosynthesis of 7S,14S-diHDHA. In addition, macrophages can express both h5-LOX and h15-LOX-1, so in the simplest terms, macrophages may be able to synthesize 7S,14S-diHDHA without participation of other cell types for transcellular biosynthesis. This is especially relevant since we did not observe platelets producing 7S,14S-diHDHA when given 7S-HDHA, therefore, the cellular milieu may affect h12-LOX activity since we observe 7S,14S-diHDHA *in vitro*. We are currently investigating these biosynthetic pathways in platelets and macrophages in more detail to identify both the pathway and the cell type that makes these oxylipins.

Second, if h15-LOX-2 is the primary enzyme involved in the *in vivo* RvD5 biosynthesis, then the role of h15-LOX-2 in human disease could be larger than expected. It is known that h15-LOX-2 converts the h5-LOX-derived pro-inflammatory oxylipins, 5S-HETE and 5,6-diHETE, into pro-resolving molecules, such as lipoxins, *in vitro* (75). With the knowledge that h15-LOX-2 also efficiently produces RvD5 from 7S-HDHA, it could be that h15-LOX-2 is important in mediating the switch between inflammation and resolution. For example, neutrophils express both h5-LOX and h15-LOX-2, so it is conceivable that neutrophils produce RvD5 independently, without the need for a transcellular synthesis mechanism. However, it should be noted that RvD5 production was observed by exposing isolated neutrophils to 17S-HpDHA (41), suggesting h5-LOX can produce RvD5 in neutrophils. This result is in contrast to our *in vitro* results and suggests that there may be a factor in the neutrophils, similar to the role of 5-lipoxygenase activating protein (FLAP), which could increase the h5-LOX activity with 17S-HDHA (76). We are currently investigating this discrepancy further by investigating neutrophils with our selective LOX inhibitors.

With respect to blood coagulation, this work shows that RvD5 has micromolar potency in inhibiting platelet activation, indicating that h15-LOX-2 may also play a role in hemostasis. RvD5 levels are reduced in blood treated with anticoagulants in vitro (77) and RvD5 formation does not occur during initial platelet activation, however, it is produced in later stages of clot progression (77). Given our result that RvD5 reduces platelet activation, the production of RvD5 could be a signal to diminish the clot size. Given that platelets do not produce h15-LOX-2 and thus cannot make RvD5, these results suggest that RvD5 is formed by h15-LOX-2 in macrophages and/or neutrophils, which migrate into the area of late-stage clots and increase their RvD5 production by increasing the h15-LOX-2 expression, and not that of h15-LOX-1 (4). We are currently investigating the role of h15-LOX-2 in more detail with our specific/potent h15-LOX-2 inhibitors (78) with the hope of understanding the correct biosynthesis pathway of RvD5 and hence its role in coagulation and atherosclerotic disorders.

## CONCLUSION

7S,14S-diHDHA is synthesized in vitro by the sequential reactions of h5-LOX and h12-LOX with DHA. However, we have also discovered a novel alternative biosynthetic pathway for the production of 7S,14S-diHDHA, which involves h15-LOX-1. The alcohol on 7S-HDHA changes the binding position of the oxylipin in the active site, thus altering the product specificity. This non-canonical result has far-reaching ramifications. It first indicates a possible alternative biosynthetic pathway for 7S,14S-diHDHA, but more importantly, the result indicates that our knowledge of LOX reactivities with fatty acids, may not extend to oxylipins, and therefore further study is needed to determine the exact biosynthetic pathways for each oxylipin, such as lipoxins, protectins and maresins. Importantly, these results suggest that LOX-products observed during *in-vivo* studies may originate from non-canonical biosynthetic routes involving previously overlooked cell types.

In addition, this result suggests that the biosynthesis of 7S,17S-diHDHA (RvD5) is achieved with h15-LOX-2 and thus its role in human disease may be more important than previously suspected. These results are especially relevant since we have discovered that both 7S,14S-diHDHA and 7S,17S-diHDHA (RvD5) are micromolar, anti-aggregation effector molecules that could possibly be synthesized directly from macrophages or neutrophils, respectively, without the need for a transcellular mechanism of biosynthesis, suggesting that both 7S,14S-diHDHA and 7S,17S-diHDHA (RvD5) may be more intimately involved in stopping the aggregation process in clot resolution than previously suspected. Thus, targeting the kinetics of platelet aggregation or clot resolution at sites of inflammation by inhibiting the biosynthesis of either 7S,14S-diHDHA and 7S,17S-diHDHA (RvD5) may represent a new therapeutic target for future development.

### Accession ID’s

P18054 = h12-LOX

O15296 = h15-LOX-2

P09917 = h5-LOX

P16050 = h15-LOX-1

## Conflict of interest

The authors declare no conflict of interest.

The content is solely the responsibility of the authors and does not necessarily represent the official views of the National Institutes of Health

## Author contributions

SP, BT, CK, MJ, MH and TH designed the experiments. SP, BT, CK and OA, SC carried out experiments. SP wrote the first draft of the manuscript, edited by TH, MH, MJ.

## Acknowledgements

### FUNDING

NIH R01 GM105671 (MH and TRH), NIH R01 HL11405 (MH and TRH), NIH R35 GM131835 (MH and TRH) and NIH K99HL136784 (BET).

